# RTTN moonlights beyond the centrosome to control ribosome biogenesis and tRNA modification in human brain organoids

**DOI:** 10.64898/2026.08.13.744412

**Authors:** A Hadar, K Draganova, V Iyer, B Bhattacharya, Y Komemy, AI Ponce-Arias, L Otikovs, I Vaknin, N Dezorella, L Wilk, A Lilja, S Doroshev, T Olender, M Danan-Gotthold, J Fu, J Merl-Pham, E Rusha, R Gabarró-Solanas, A Flatley, H Zitzelsberger, R Feederle, SM Hauck, S Schwartz, M Götz, O Reiner

## Abstract

RTTN (rotatin) is a centrosomal protein mutated in severe malformations of cortical development, yet how its dysfunction disrupts human corticogenesis has remained unclear. Here, we show that RTTN has an unrecognized function at the core of the translation machinery. Using human telencephalic and hippocampal organoids carrying distinct *RTTN* alleles, together with single-cell and bulk transcriptomics, polysome profiling, and tRNA pseudouridine sequencing, we find that *RTTN* is enriched in cycling first-trimester neural progenitors and physically associates with ribosome-biogenesis and RNA-processing factors. *RTTN* mutations impair rRNA biogenesis and polysome assembly, reduce cytoplasmic ribosome density and nascent protein synthesis, and remodel the tRNA pseudouridylation landscape through both a PUS7L-dependent variable-arm signature and a broader *RTTN*-specific defect. These translational deficits are accompanied by prolonged mitosis, reduced entry into S-phase, and impaired interkinetic nuclear migration in mutant progenitors. Our findings redefine RTTN as a regulator of ribosome homeostasis and mRNA translation and implicate defective translational capacity as a driver of *RTTN*-associated microcephaly.

**Graphical Abstract:** RTTN sustains ribosome and tRNA homeostasis in human neural progenitors; its mutation disrupts mRNA translation, stalling progenitor proliferation and interkinetic nuclear migration, and driving cortical malformation and growth failure.

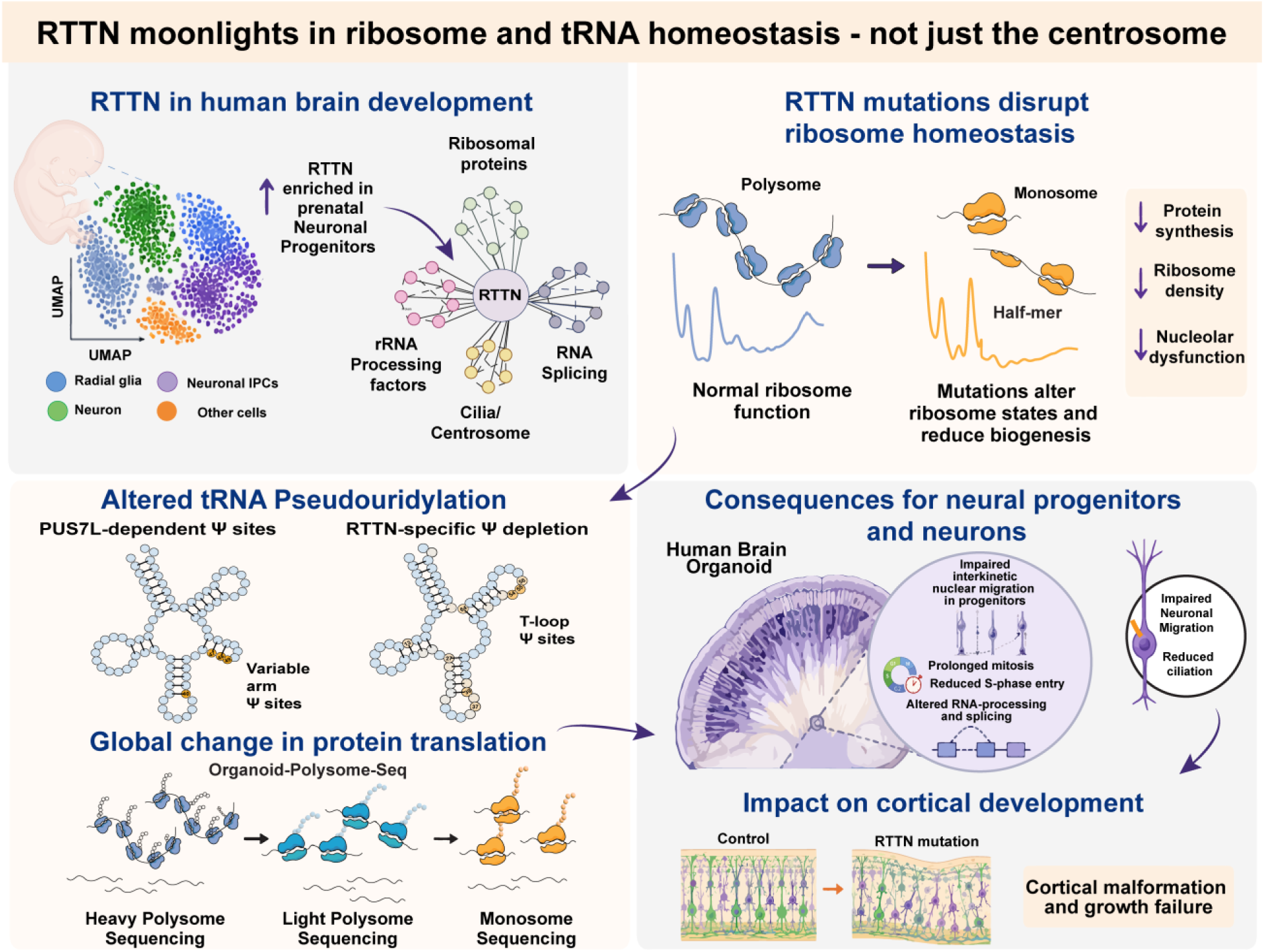

## Introduction

Congenital malformations of cortical development (MCDs), including primary microcephaly and polymicrogyria, are genetically heterogeneous but repeatedly converge on core cellular processes in neural stem and progenitor cells, including centrosome/centriole biogenesis, primary cilium signaling, mitotic progression, and apical-basal epithelial organization^1^. Pathogenic variants in *RTTN* (encoding the centrosomal protein rotatin) were first linked to polymicrogyria and disordered cortical organization, connecting centrosome/cilium dysfunction to abnormal human corticogenesis^2–4^. Subsequent clinical-genetic studies expanded the *RTTN* spectrum to include primary microcephaly and primordial dwarfism, suggesting that *RTTN* dosage and allele-specific effects can impair brain growth across developmental contexts^2–4^. These observations position *RTTN* among core developmental cell-biology genes in which relatively partial perturbations of fundamental cellular processes can produce severe neuroanatomical consequences.

Mechanistically, RTTN is a conserved centrosome-associated protein required for centriole elongation and maturation. RTTN localizes to procentrioles, interacts with canonical centriole-assembly regulators, and is necessary for the formation of full-length centrioles and the recruitment of distal centriolar components, with downstream implications for centrosome function and ciliogenesis^5^. These requirements align with patient phenotypes that overlap with ciliopathies and other centrosomopathies, in which progenitor pool maintenance and the balance between symmetric/asymmetric division are especially sensitive to centrosome integrity. Consistent with this framework, recent studies have demonstrated that a Taybi-Linder syndrome-related *RTTN* variant impairs neural rosette formation, delays apicobasal polarization, and induces mitotic/cell-cycle abnormalities in human cortical organoids, highlighting a tractable human system in which *RTTN* dysfunction can be linked to early tissue architecture^6^. Despite evidence that RTTN disruption compromises neuroepithelial organization and cell-cycle progression, the downstream molecular programs that amplify these defects into tissue-level failure (e.g., impaired growth and reduced neurogenic output) remain unknown.

Human brain organoids are well-suited for the mechanistic dissection of *RTTN*-associated brain disease because they capture quantifiable features of proliferation within a polarized neuroepithelium, interkinetic nuclear migration, and staged neurogenesis, while enabling controlled comparisons between patient-derived and gene-edited isogenic lines^7–9^. Here, using human brain organoids and single-cell, multi-omic analyses, we identify an unanticipated role for RTTN in maintaining ribosome homeostasis and translational capacity during human corticogenesis. We show that RTTN is enriched in proliferative neural progenitors and forms an interactome associated with ribosomal and RNA-processing factors. *RTTN* mutations disrupt rRNA-related pathways, polysome engagement, and nascent protein synthesis, accompanied by impaired progenitor proliferation and neuronal development. Together, our findings uncover an unexpected role for RTTN beyond its established centrosomal functions, demonstrating that RTTN is also required to maintain ribosome homeostasis and translational capacity during human corticogenesis.

## Results

To determine when and where *RTTN* is expressed during human neurodevelopment, we analyzed transcriptomic resources spanning prenatal and postnatal periods of human brain development^10^. *RTTN* mRNA levels were significantly higher at prenatal stages than at postnatal ages in BrainSpan (https://www.brainspan.org/; Extended Data Fig. 1a), consistent with its predominant role during early corticogenesis. Analysis of single-cell RNA-seq from the first-trimester developing human brain (PCW 5-14^11^) further revealed that *RTTN* is expressed in a lineage- and stage-specific manner during neurogenesis. Specifically, *RTTN* expression was highest in neuronal intermediate progenitor cells (IPCs) compared with radial glia, neuroblasts, and mature neurons (Fig. 1a, Extended Data Fig. 1b,c). Along with the neurogenic trajectory, *RTTN* expression peaked in IPCs and declined during neuronal differentiation, while remaining relatively stable in radial glia and glioblasts. *RTTN* expression was also reduced in cells associated with the oligodendrocyte lineage, consistent with its enrichment in proliferative neural progenitor populations. Developmental heatmap analysis further showed consistent *RTTN* enrichment in IPCs across developmental stages (Fig. 1b). Within radial glia, *RTTN* expression was broadly distributed across brain regions (Extended Data Fig. 1b), suggesting its deployment across multiple neuroepithelial compartments. Consistent with a role in cell proliferation, *RTTN* expression was highest in cycling cells, particularly in S and G2/M phases, and reduced in non-cycling or post-mitotic populations (Fig. 1c, Extended Data Fig. 1d).

**Fig. 1:**
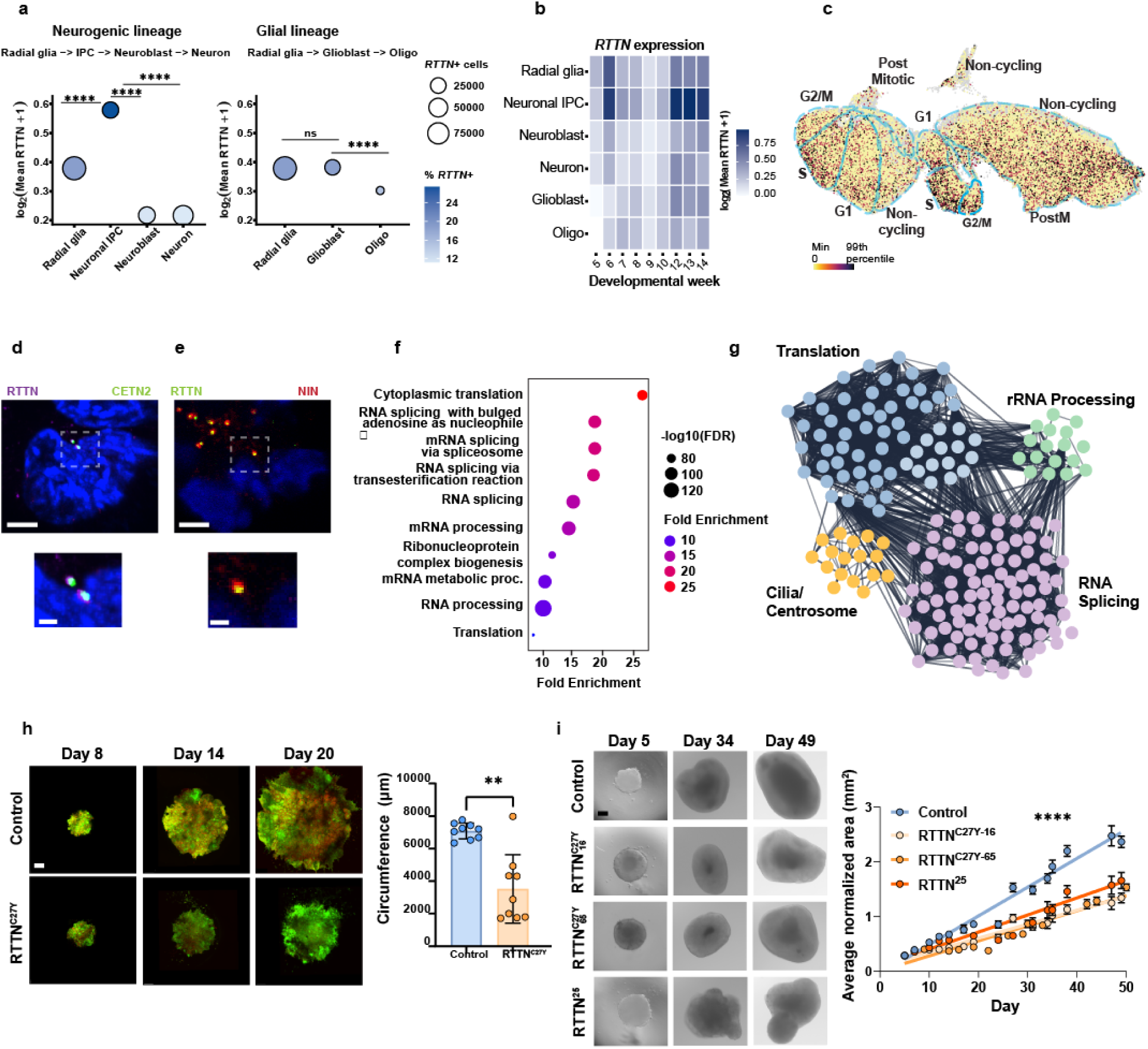
RTTN is enriched in prenatal neural progenitors and associates with translation and rRNA-processing factors. **a,** *RTTN* expression across neurogenic (Radial glia → Neuronal IPC → Neuroblast → Neuron) and glial (Radial glia → Glioblast → Oligo) developmental trajectories^11^ (Braun et al., Science 2023). Dot size indicates the percentage of *RTTN*-positive cells, while color intensity represents mean *RTTN* expression calculated as log2(mean *RTTN* + 1). *RTTN* expression was significantly enriched in neuronal intermediate progenitor cells (IPCs) compared with radial glia, neuroblasts, and neurons. In the glial lineage, *RTTN* expression was maintained between radial glia and glioblast populations but significantly decreased in oligodendrocyte lineage cells. Statistical significance was assessed using Wilcoxon rank-sum tests with Benjamini-Hochberg correction. ns, not significant; ****P < 0.0001. **b,** Heatmap showing *RTTN* expression across developmental weeks and neural cell populations. Color intensity represents mean *RTTN* expression calculated as log2(mean *RTTN* + 1). **c,** UMAP projection of single-cell RNA-seq from the first-trimester developing human brain of forebrain (*EMX1-*positive) radial glia annotated by cell-cycle phase (G1, S, G2/M, post-mitotic (PostM), and non-cycling), showing the distribution of *RTTN*-expressing cells across proliferative states with enrichment in G2/M and S phase. **d,** RTTN-HaloTag knock-in (magenta) hESCs. Immunofluorescence staining reveals that RTTN localizes to the centrosome, as indicated by colocalization with CETN2 (centrin 2, green), a structural centrosomal protein. Scale bar, 4 µm; zoom scale bar, 1 µm. **e,** Immunostaining of RTTN in d15 neural progenitor cells shows that the protein localizes to centrosomes labeled with Ninein (NIN), scale bar, 4 µm; zoom scale bar, 1 µm. **f,** Gene Ontology biological process enrichment analysis of RTTN-interacting proteins. Bubble size indicates gene count, and color indicates FDR (as shown). **g,** Protein-protein interaction network of the RTTN interactome (derived from RTTN immunoprecipitation followed by mass spectrometry and filtered for enriched interactors), visualized in Cytoscape/STRING. Nodes represent RTTN-enriched interacting proteins, and edges represent known or predicted functional associations. Major functional modules are annotated (translation, rRNA processing, RNA splicing, and Cilia/Centrosome). Statistical significance was assessed using Welch’s t-test, P < 0.05, s_0_ = 0.6. **h,** Representative time-course images of on-chip brain organoids (green-LYN-GFP, red-H2B mCherry) comparing Control and RTTN^C27Y^ (days 8, 14, and 20) show limited growth in mutant organoids. Scale bar, 200 µm. Quantification of organoid size from h (circumference, µm) for Control (n=9) and *RTTN*^C27Y^ (n=9); each dot represents one organoid, three biological batches; Two-tailed Mann-Whitney U test, P = 0.0040. **i**, Representative images of hippocampal organoids, and growth curves of hippocampal organoids (average normalized area over time) comparing Control with *RTTN* mutant lines (*RTTN*^C27Y^ lines and mutated *RTTN*^25^lines as labeled); points show mean with error bars, and fitted trends are indicated. Linear regression analysis revealed significant differences in slopes between groups, indicating slower growth of *RTTN* mutant organoids (F(3, 521) = 40.69, P < 0.0001). Individual regressions were significant for all groups (P < 0.0001, R² = 0.58–0.81). Data represent measurements from n = 4-17 organoids per group per day, two biological batches.

To visualize endogenous RTTN protein, we generated a HaloTag knock-in human embryonic stem cell (hESC) line. HaloTag-RTTN localized to the centrosome, as demonstrated by colocalization with the centrosomal marker CETN2 (Fig. 1d), consistent with previous studies^6, 12, 13^. Immunostaining of RTTN in Day 15 neural progenitor cells (NPCs) generated by directed neuronal differentiation from iPSCs further demonstrated that the RTTN protein localizes to centrosomes marked by Ninein (NIN) (Fig. 1e).

We next sought to define the RTTN protein interaction landscape by immunoprecipitating endogenous RTTN from neural progenitor lysates and analyzing the immune complexes by label-free liquid chromatography tandem mass spectrometry (MS). Surprisingly, Gene Ontology analysis indicated enrichment of translation- and ribosome-associated pathways among RTTN interactors (Fig. 1f), identifying RNA metabolism and translational control as key functional axes linked to RTTN. Network analysis of RTTN-enriched interactors revealed distinct functional modules, prominently including proteins involved in translation, rRNA processing, RNA splicing, and centrosome/ciliary biology (Fig. 1g; Extended Data Fig. 1g). These results point to an unexpected functional role of RTTN in translational regulation and RNA metabolism.

We then assessed the functional consequences of distinct *RTTN* mutations using hESC-derived brain organoids (Extended data Fig. 2). Specifically, we first generated *RTTN*^C27Y^ brain-on-chip organoids, following a previously published protocol^7^. *RTTN*^C27Y^ organoids exhibited reduced growth relative to wild-type (WT) controls throughout the culture period (Fig. 1h, Extended data Fig. 2,3). Similarly, in hippocampal organoids, *RTTN* mutant lines showed impaired growth dynamics relative to controls (Fig. 1i). These data establish that *RTTN* mutations impair early brain organoid growth, consistent with impaired proliferative capacity, and, given the interactome findings, suggest translational homeostasis as a downstream vulnerability during corticogenesis.

To test whether *RTTN* mutations alter translational output, we quantified nascent protein synthesis by measuring O-propargyl-puromycin (OPP) incorporation^14^. In hESCs, *RTTN*^C27Y^ did not measurably affect OPP incorporation relative to control cells (Fig. 2a, Extended data Fig. 2), indicating that basal protein synthesis is not impaired in this undifferentiated state. We next examined *RTTN* mutant organoids in a neurodevelopmentally relevant context. Hippocampal organoids day 84 were dissociated and acutely labeled with OPP to measure nascent polypeptide production at the single-cell level (Fig. 2b). Cells derived from *RTTN*-mutant hippocampal organoids derived from *RTTN*-mutant hESCs, including *RTTN*^C27Y^ and two additional *RTTN* mutant lines, *RTTN*^26^, and *RTTN*^10^, showed reduced OPP signal compared with two independent control lines (Fig. 2c), indicating that RTTN-dependent impairment of protein synthesis emerges during neural differentiation and/or within organoid tissue contexts. Moreover, this reduction in nascent protein synthesis was generalized across *RTTN* alleles and organoid platforms, including iPSC-derived *RTTN*^R985G^ organoids, which exhibited reduced OPP incorporation relative to control iPSC-derived organoids (Fig. 2d). Together, these findings establish that *RTTN* mutations reduce nascent protein synthesis in hippocampal organoids, supporting a role for RTTN in maintaining translational capacity during human neurodevelopment.

**Fig. 2:**
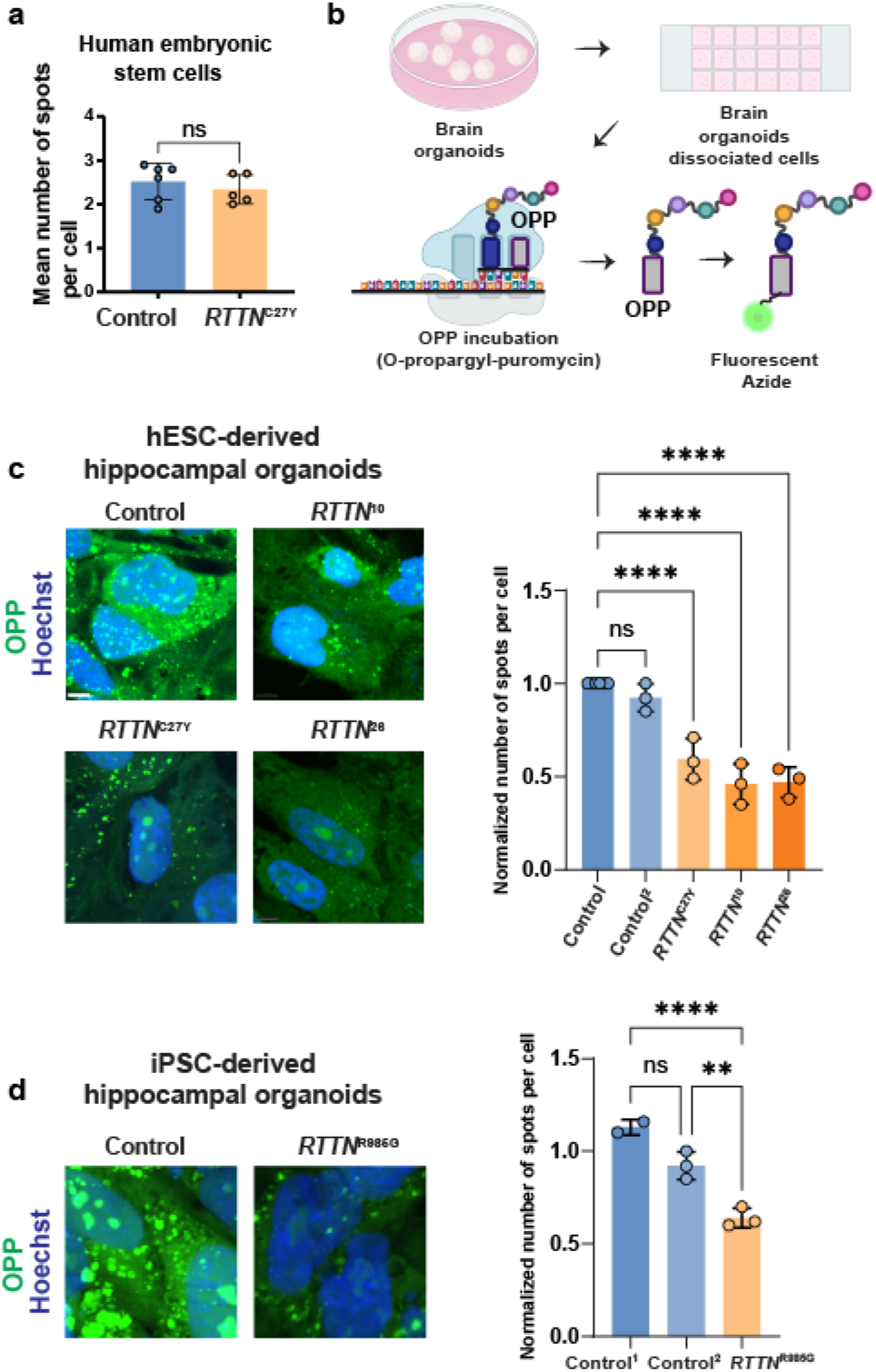
*RTTN* mutations reduce nascent protein synthesis in hippocampal organoid-derived cells and iPSC-derived hippocampal organoids. **a**, Translation rate in human embryonic stem cells (hESCs) carrying *RTTN*^C27Y^ and control hESCs measured by incorporation of the puromycin analog O-propargyl-puromycin (OPP). Quantification shows the average number of OPP-positive spots per cell; no significant difference was detected between control (n = 6 images, 1615 cells) and *RTTN*^C27Y^ (n = 5 images, 1534 cells) (P= 0.4481, Mann-Whitney test). The images shown are from a single representative experiment; the quantifications are normalized across three independent biological batches. **b,** Schematic of the OPP incorporation assay performed on hippocampal organoids. Organoids were dissociated into single cells and incubated with OPP for 1 h, followed by click-chemistry detection using Alexa Fluor 488 Picolyl Azide to label nascent polypeptides. **c**, Representative fluorescence images of cells dissociated from hESC-derived hippocampal organoids labeled with Hoechst (nuclei, blue) and OPP (nascent polypeptides, green), Shown are Control, Control^2^, *RTTN*^C27Y^, *RTTN*^26^, and mutated *RTTN*^10^. Control², hESC control line edited with HaloTag-tagged RTTN. Control n=6, Control^2^ n=3, *RTTN*^C27Y^ n=3, *RTTN*^26^ n=3, *RTTN*^10^ n=3 independent experiments; each dot is the average of an independent experiment. One-way ANOVA, p<0.0001, F=44.53. Holm-Sidak’s multiple comparisons revealed no difference between control and control^2^ (adjusted P = 0.1464), with significant differences between control and all *RTTN* mutant groups (adjusted P < 0.0001). Scale bar, 5 µm. **d,** Representative fluorescence images of iPSC-derived hippocampal organoids labeled with Hoechst (blue) and OPP (green) from iPSC control^1^, Control^2^, and *RTTN*^R985G^ (pArg985Gly) lines. Quantification (right) shows the normalized mean number of OPP-positive spots per cell; (Control n=2, Control^2^n=3, RTTN^R985G^ n=3, independent experiments, one-way ANOVA p< 0.0001) with Holm-Šídák’s multiple comparisons test showed no difference between control^1^ and control^2^ (P = 0.516), but significant differences involving *RTTN*^R985G^ (P < 0.0001 and P = 0.0015). Each dot represents an independent experiment. Scale bar, 5 µm.

To further elucidate the mechanistic basis of reduced translational capacity, we performed polysome profiling and polysome fractionation coupled with RNA-seq (polysome-seq) in telencephalon organoids to quantify ribosome-state distributions and transcript-specific ribosome engagement (Fig. 3, Extended Data Fig. 3). Under basal conditions, *RTTN*^C27Y^ organoids showed a clear shift in the polysome distribution compared with controls (Fig. 3b,c), with a significant decrease in the polysome-to-monosome (P/M) ratio at both day 60 and day 30 (Fig. 3c,d), indicating that impaired polysome formation is present across developmental stages of *RTTN*^C27Y^ organoids. Additionally, *RTTN*^C27Y^ organoids exhibited altered ribosomal subunit balance. The 40S/60S ratio measured for *RTTN*^C27Y^ organoids under native conditions was significantly smaller relative to control (Fig. 3e), suggesting disruption of ribosome homeostasis and/or subunit joining due to *RTTN* mutations. Importantly, EDTA-mediated dissociation of ribosomes abolished the genotype-dependent difference in the 40S/60S ratio between *RTTN*^C27Y^ and control telencephalon organoids (Fig. 3f), demonstrating that this imbalance depends on genotype-regulated ribosome assembly state rather than a potential technical variation. Together, these data show that *RTTN* mutation reshapes ribosome-state distributions in telencephalon organoids, with both reduced polysome engagement and altered subunit homeostasis.

**Fig. 3.**
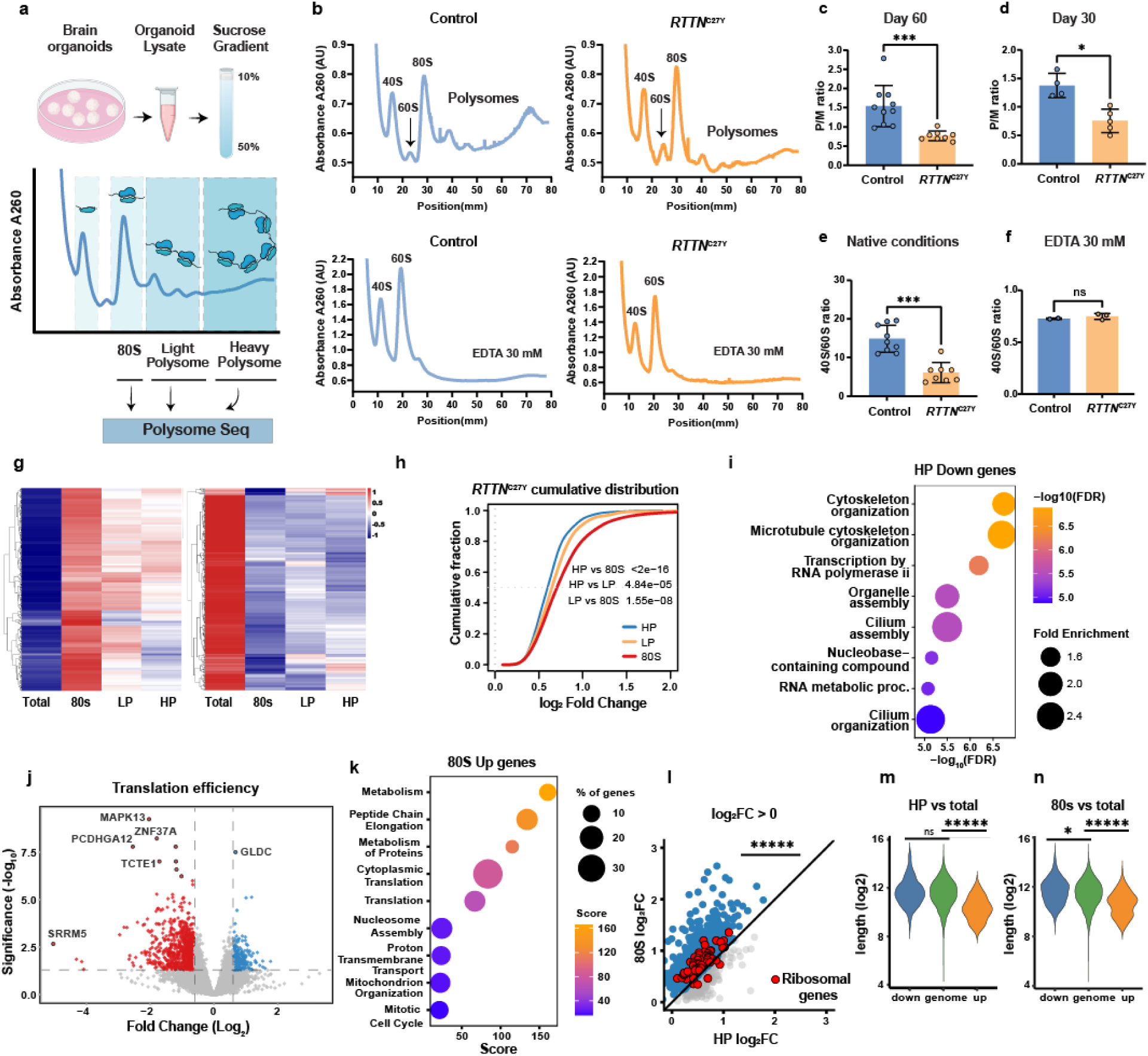
*RTTN* mutation alters ribosome homeostasis and translational efficiency in human telencephalon organoids. **a**, Schematic of the experimental workflow for polysome profiling and polysome-seq analysis. **b**, Representative absorbance 260 polysome profiles from control and *RTTN*^C27Y^ telencephalon organoids fractionated on sucrose gradients. Peaks corresponding to 40S, 60S, 80S (monosome), and polysomes are indicated. Lower traces show profiles following EDTA (30 mM) treatment, which dissociates ribosomes into free subunits. **c**, Quantification of the areas under the curve (AUC) of polysome-to-monosome (P/M) ratio in day 60 telencephalon organoids, showing reduced P/M in *RTTN*^C27Y^ (Each dot represents a single gradient run (n = 8), performed on lysates pooled from at least 10 individual organoids.) compared with control (n=10, gradient run). Two-tailed Mann-Whitney test, U = 2, P = 0.0002. **d**, Quantification of the P/M ratio in day 30 telencephalon organoids in *RTTN*^C27Y^ (n=5, gradient run) compared with control (n=4, gradient run) Two-tailed Mann-Whitney test U = 0, P = 0.0159. **e**, Quantification of 40s/60s ratio under native conditions in telencephalon organoids, expressed as the 40S/60S ratio, indicating altered subunit homeostasis in *RTTN*^C27Y^ in *RTTN*^C27Y^ (n=8, gradient run) compared with control (n=9, gradient run). Two-tailed Mann-Whitney test, U = 2, P = 0.0003. **f**, 40S/60S ratio after EDTA (30 mM) treatment (ns), indicating that the genotype-dependent difference in subunit ratio is lost after ribosome dissociation in *RTTN*^C27Y^ (n=3, gradient run) compared with control (n=2, gradient run). Two-tailed Mann-Whitney test U = 2, P = 0.8. **g**, Heatmaps showing scaled gene expression across Total RNA, 80S, light polysome (LP), and heavy polysome (HP) fractions in *RTTN*^C27Y^ organoids. Rows represent genes, columns represent fractions, and hierarchical clustering is applied; the color scale indicates relative expression. h, Cumulative distribution of log₂ fold change (normalized to 1) for genes in HP, LP, and 80S fractions, showing a progressive shift from HP to LP to 80S. Statistical comparisons between groups were performed using the Kolmogorov-Smirnov test (P-values indicated in the figure). **i**, Gene Ontology enrichment analysis of downregulated genes in the HP fraction, showing terms related to cytoskeleton organization, cilium assembly, and RNA metabolism. Dot size represents fold enrichment, and color indicates -log₁₀(FDR). **j**, Volcano plot showing differential translational efficiency (polysome-seq) between control and *RTTN*^C27Y^ telencephalon organoids (log2 fold change versus -log10 adjusted P value). Significantly altered transcripts are highlighted (color coding as shown). Adjusted P value<0.05, log2 fold change cutoff=0.6, red dots indicate decreased TE, blue dots indicate increased TE in mutant relative to control. **k,** Gene Ontology enrichment analysis of upregulated genes in the 80S fraction, highlighting enrichment of ribosomal pathways, cell cycle-related processes, and mitochondrial organization. Dot size represents the percentage of genes, and color indicates the enrichment score. **l,** Scatterplots showing log₂ fold change (*RTTN*/control) of transcript abundance in polysome profile fractions normalized to total RNA. Each point represents a gene, with ribosomal protein genes highlighted in red. Axes compare enrichment between fractions (80S, HP), and the diagonal line denotes equal fold change. Positive log₂FC indicates increased representation in *RTTN*. Ribosomal protein transcripts cluster tightly and show a pronounced shift toward the 80S fraction. Pairwise comparisons were performed using paired Wilcoxon signed-rank tests (P<0.00001). **m,n,** Distribution of transcript lengths (log₂) for downregulated, genome-wide, and upregulated genes in HP (m) and 80S (n) fractions. Statistical significance was assessed as indicated (ns, not significant; *P < 0.05; ***P < 0.0001).

To determine whether these global changes translate into transcript-specific effects, we performed polysome-seq and analyzed RNA distribution across total RNA, 80S (monosomes), light polysome (LP), and heavy polysome (HP) fractions (Fig. 3g). *RTTN*^C27Y^ telencephalon organoids exhibited widespread redistribution of transcripts across fractions, with cumulative distribution analysis revealing a shift from HP toward LP and 80S fractions (Fig. 3h), consistent with reduced translational engagement. Gene Ontology analysis of transcripts depleted from the HP fraction highlighted cellular processes related to cytoskeleton organization, cilium assembly, and RNA metabolism (Fig. 3i), while transcripts enriched in the 80S fraction were associated with ribosomal and cell cycle pathways (Fig. 3k). Consistent with these findings, differential translational efficiency analysis identified significantly altered transcripts in *RTTN*^C27Y^ telencephalon organoids (Fig. 3j), and fraction-resolved comparisons showed a pronounced redistribution of ribosomal protein mRNAs toward the 80S fraction (Fig. 3l), indicative of stalled or non-productive ribosome engagement. Furthermore, transcript length analysis revealed a significant bias toward shorter transcripts among upregulated genes, with transcripts enriched in the HP and 80S fractions significantly shorter than those in the total transcriptome (Fig. 3m,n), indicating that *RTTN* mutation preferentially affects transcript engagement with ribosomes by transcript length. Codon-level analysis of fraction-specific transcript redistribution revealed a subset of codons with highly significant depletion (fold change > 2, adjusted P < 1 × 10⁻^6^) across ribosome fractions, with distinct patterns across 80S, LP, and HP fractions (Supplementary Table 1). Notably, ATA and TTA codons were depleted across all fractions, suggesting impaired ribosome engagement that extends from initiation through elongation. In contrast, TCA codons were selectively depleted from the LP and HP fractions but not from the 80S fraction, consistent with preserved initiation but reduced elongation progression. A third group of codons (AAA, AGA, CAA, and GAA) showed depletion restricted to the LP fraction, suggesting an early elongation delay prior to efficient accumulation in heavier polysomes. Together, these graded patterns across fractions point to codon-specific differences in translational efficiency, with effects most prominent for A-ending codons, suggesting non-uniform perturbation of decoding dynamics rather than a global translational shutdown.

To determine whether *RTTN* mutations alter ribosome assembly states in a differently regionalized brain context, we performed sucrose-gradient polysome profiling in hippocampal organoids derived from control and *RTTN*^26^, *RTTN*^10^, and *RTTN*^C27Y^ mutant hESC lines. Compared with controls, *RTTN*-mutant hippocampal organoids displayed prominent half-mer-like features in the polysome region (Fig. 4a), and quantification confirmed a significant increase in the half-mer fraction (Fig. 4b), consistent with impaired 60S subunit joining. Moreover, *RTTN* mutants exhibited significant shifts in ribosome-state ratios, including altered 40S/60S and 80S/60S (Fig. 4c,d), although the 80S/40S ratio did not reach statistical significance (Fig. 4e). Together, these profiles indicate disrupted ribosome assembly and subunit balance. Consistent with this, TapeStation analysis of purified total RNA revealed an increased 28S/18S ratio in *RTTN* mutant samples, suggesting altered steady-state rRNA homeostasis and an imbalance in 40S/60S ribosomal subunits (Fig 4f).

**Figure 4.**
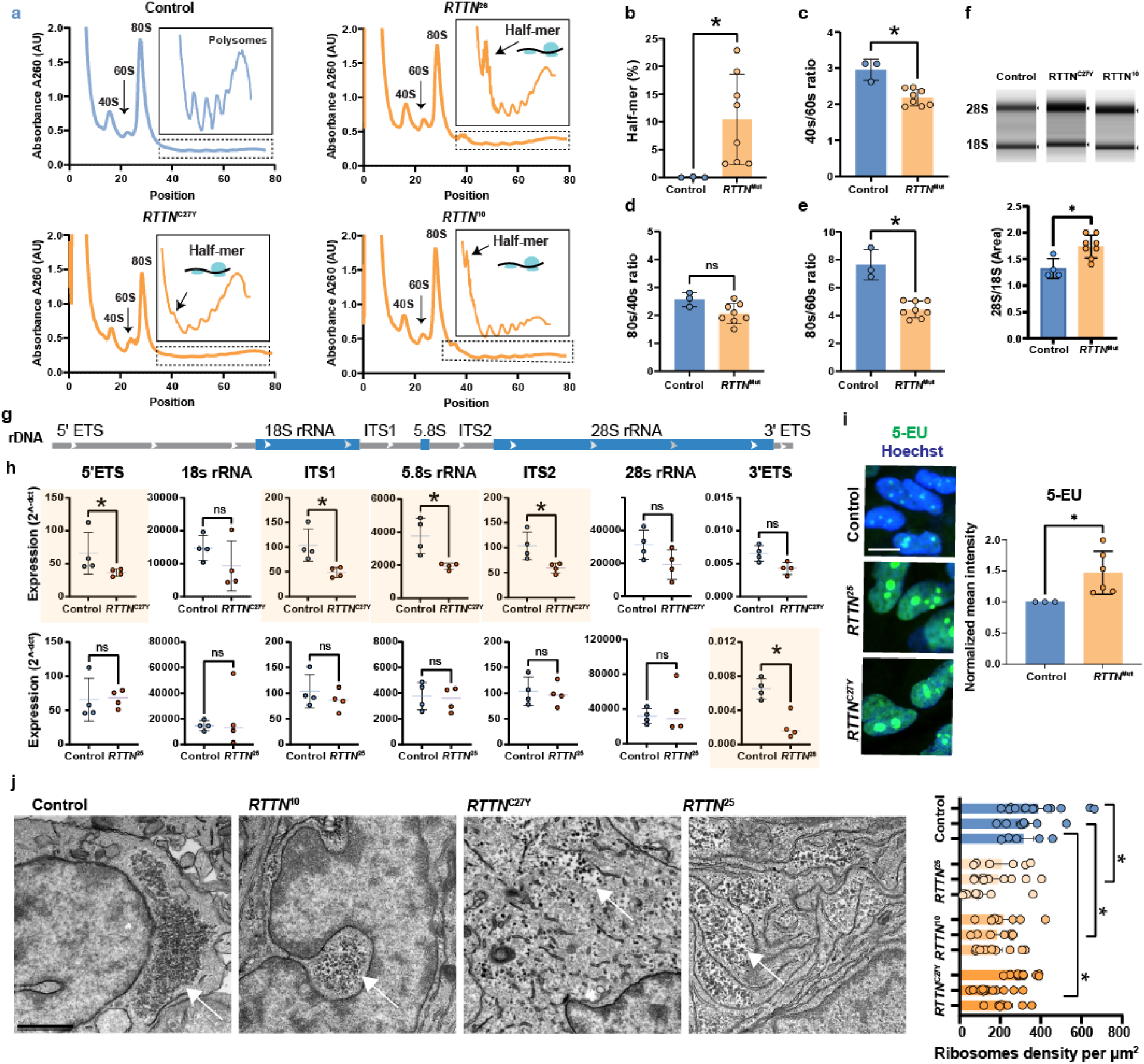
*RTTN* mutations perturb the balance of ribosomal states and rRNA biogenesis in hippocampal organoids. **a**, Representative A260 polysome profiles from day 50 control hippocampal organoids and *RTTN*-mutant organoids (*RTTN*^26^, *RTTN*^10^, and *RTTN*^C27Y^). Lysates were separated on sucrose gradients, and absorbance was monitored across fractions. Peaks corresponding to 40S, 60S, 80S (monosome), and polysomes are indicated. Insets highlight half-mer-like features in *RTTN*-mutant profiles. **b**, Quantification of half-mer (%) normalized to 80s peak comparing control (each dot represents a single gradient run (n = 3), performed on independent lysates pooled from at least 10 individual organoids and mutated *RTTN* (pooled across *RTTN*^26^, *RTTN*^10^, and *RTTN*^C27Y^), n = 8 independent gradient runs, 8 organoids in each gradient. Two-tailed Mann-Whitney test, U = 0, P = 0.0121. **c-e**, Quantification of ribosome-state area-under-curve ratios derived from polysome profiles: 40S/60S Two-tailed Mann-Whitney test U = 0, P = 0.0121.(c), 80S/40S Two-tailed Mann-Whitney test U = 3, P = 0.0848(d), and 80S/60S Two-tailed Mann-Whitney test U = 0, P = 0.0121 (e) for control versus mutated *RTTN*. Each dot represents an independent measurement; bars show group means with error bars as plotted. **f**, Representative TapeStation electropherograms showing 28S and 18S rRNA peaks from Control (n=4), and *RTTN*^Mut^ *(RTTN*^C27Y^ (n=4) and *RTTN*^10^ (n=4) grouped together) hippocampal organoids purified RNA (top). Quantification of the 28S/18S rRNA ratio (area under the peaks) revealed a significant increase in *RTTN* mutant samples compared with controls (bottom). Each dot represents an individual sample; bars indicate mean ± SD. Statistical significance was determined using a two-tailed Mann-Whitney test. P=0.0182. g. Schematic of the human rDNA transcription unit indicating regions interrogated by RT-qPCR: 5′ ETS, 18S rRNA, ITS1, 5.8S rRNA, ITS2, 28S rRNA, and 3′ ETS. **h**, RT-qPCR of rDNA regions in control (n=4), *RTTN*^C27Y^ (n = 4) and *RTTN*^25^ (n=4) samples. Each dot represents one biological replicate comprising at least seven independent hippocampal organoids. Data were normalized to the UBC reference gene using the 2^^−ΔCt^ method. Analysis of pre-rRNA and mature rRNA species revealed significant alterations in *RTTN*^C27Y^ cells compared with Control cells. ITS1, ITS2, 5′ ETS, and 5.8S rRNA levels were significantly reduced in *RTTN*^C27Y^ cells (two-tailed Mann-Whitney U test, U = 16, p = 0.0286 for each comparison; n = 4 per group). 3′ ETS levels also showed a pronounced reduction in *RTTN*^C27Y^ cells, although this difference did not reach statistical significance (U = 15, p = 0.0571). No significant differences were observed for 5S rRNA (p = 0.3429), 18S rRNA (p = 0.3429), or 28S rRNA (p = 0.2000). 3′ ETS levels were significantly reduced in *RTTN*^25^ compared with Control cells (U = 16, p = 0.0286), two-tailed Mann-Whitney U tests, whereas no significant differences were detected for the other analyzed rRNA regions.; ns, not significant. **i,** Representative images of hippocampal organoids with 1 hour incubation of 5-ethynyl uridine (5-EU) incorporation (nascent RNA; green) with Hoechst nuclear staining (blue) in control (n=3), *RTTN*^C27Y^ (n=3), and mutated *RTTN*^25^ (n=3), scale bar 15 µm. Right: quantification of normalized mean 5-EU intensity; mutations grouped together (Two-tailed Mann-Whitney test, P=0.0238); 3 independent experiments; each dot is the average of an independent experiment. **j,** Representative transmission electron microscopy (TEM) images of cytoplasmic regions from control and *RTTN*-mutant organoids (day 90) (*RTTN*^C27Y^, *RTTN^10^*, *RTTN*^25^). Ribosomes are visible as electron-dense particles in the cytoplasm. Scale bar, 1 µm. Quantification of ribosome density from TEM images. For each condition, three independent organoids were analyzed, and 5–14 micrographs per organoid were quantified using the ParticleSizer plugin in Fiji (NanoDefine). Ribosome density was calculated as the number of ribosomes per µm² within defined regions of interest (ROIs). Each dot represents one analyzed ROI; bars indicate mean ± s.e.m. Statistical significance was assessed using a nested one-way ANOVA followed by Holm-Šídák’s multiple-comparisons test: Control versus *RTTN*^25,^ P=0.015; Control versus *RTTN*^10,^ P=0.024; Control versus *RTTN*^C27Y^, P=0.037.

Altered ribosome states can arise from defects in ribosome biogenesis^15–17^; we therefore assessed pre-rRNA and rRNA processing intermediates using RT-qPCR across the rDNA transcription unit (Fig. 4g). *RTTN*^C27Y^ telencephalon organoids showed significantly reduced levels of multiple ITS- and ETS-containing pre-rRNA processing intermediates (Fig. 4h), whereas mature 18S and 28S rRNA levels remained unchanged, suggesting impaired early pre-rRNA processing and/or reduced stability of precursor intermediates. Telencephalon organoids from a second *RTTN* mutant line (*RTTN*^25^) showed a partially overlapping but distinct pattern (Fig. 4i), indicating that defects in rRNA biogenesis may differ in extent or at the stage affected across *RTTN* mutations.

To assess nascent RNA synthesis, we performed 5-ethynyl uridine (5-EU) labeling followed by click-chemistry detection. Mutated *RTTN*^C27Y^ and *RTTN*^25^ telencephalon organoids exhibited increased 5-EU signal intensity compared with WT control (Fig. 4i), suggesting dysregulated transcriptional output, which is consistent with compensatory or perturbed nucleolar activity in the context of defective rRNA maturation. Finally, transmission electron microscopy revealed reduced cytoplasmic ribosome density in *RTTN-*mutant telencephalic organoids compared with WT controls (Fig. 4j), providing ultrastructural evidence of impaired ribosome homeostasis. Together, these data show that *RTTN* mutations disrupt ribosome assembly, perturb rRNA biogenesis, induce nucleolar stress responses, and reduce cytoplasmic ribosome abundance, providing a mechanistic link between *RTTN* dysfunction and impaired translational capacity in developing human brain tissue.

To determine whether the ribosome biogenesis defects observed in RTTN-mutant organoids extend to other RNA-processing pathways, we next investigated tRNA pseudouridylation. Because tRNA maturation and modification occur in close coordination with nucleolar RNA metabolism and are required for efficient translation, we hypothesized that disruption of ribosome homeostasis in *RTTN*-mutant organoids might also alter the tRNA pseudouridylation landscape. We therefore performed BACS-based tRNA pseudouridine sequencing (Ψ-seq) in hippocampal organoids derived from control, *RTTN*^KO25^, *RTTN*^C27Y^, and two independent *PUS7L*-knockout lines (Extended Data Fig. 4a).

Transcriptomic analysis identified *PUS7L*, a tRNA pseudouridine synthase, as one of the most significantly downregulated genes in *RTTN*-mutant day 49 hippocampal organoids (Fig. 5a, Extended Data Fig. 4a), a finding validated by RT-qPCR in both hESCs, day 22 and day-34 hippocampal organoids (Fig. 5b). Given the established role of PUS7L in catalyzing pseudouridylation within the variable arm of specific tRNAs^18^, we first asked whether *RTTN* deficiency recapitulated the canonical PUS7L-dependent modification signature. Comparison of two independent *PUS7L* knockout lines (Extended Data Fig. 4b-c) with *RTTN*^KO25^ and *RTTN*^C27Y^ identified a reproducible core set of eleven shared pseudouridylation sites, primarily located within Leu and Ser tRNAs (Fig. 5c,d, Extended Data Fig. 4d). Structural mapping localized these shared sites to the canonical variable-arm positions 46-49 (Fig. 5e), consistent with the known substrate specificity of PUS7L^18^. Additional shared new identified sites were at tRNA positions 39 and 40, which mapped outside the canonical variable arm. Unexpectedly, *RTTN* mutants *RTTN*^KO25^ and *RTTN*^C27Y^ exhibited a substantially broader pseudouridylation phenotype that extended beyond the shared PUS7L-dependent signature. With 59.0% and 58.4% of all candidate sites significantly altered (196/332 and 194/332, respectively), compared with only 9.3% and 8.7% in *PUS7L*^KO-7^ and *PUS7L*^KO-66^. Although the canonical PUS7L variable-arm sites were consistently depleted, the majority of *RTTN*-associated changes occurred at additional structural positions absent from the *PUS7L* knockout profile (Fig. 5c). Mapping these *RTTN*-specific sites onto a canonical tRNA secondary structure revealed a striking enrichment within the T-loop, particularly at positions 54-56, which accounted for approximately 37% of all *RTTN*-associated depleted Ψ sites (Fig. 5f). Thus, *RTTN* mutations induce both the expected PUS7L-dependent variable-arm pseudouridylation signature and an additional, previously unrecognized T-loop-associated pseudouridylation defect affecting multiple tRNA families.

**Figure 5.**
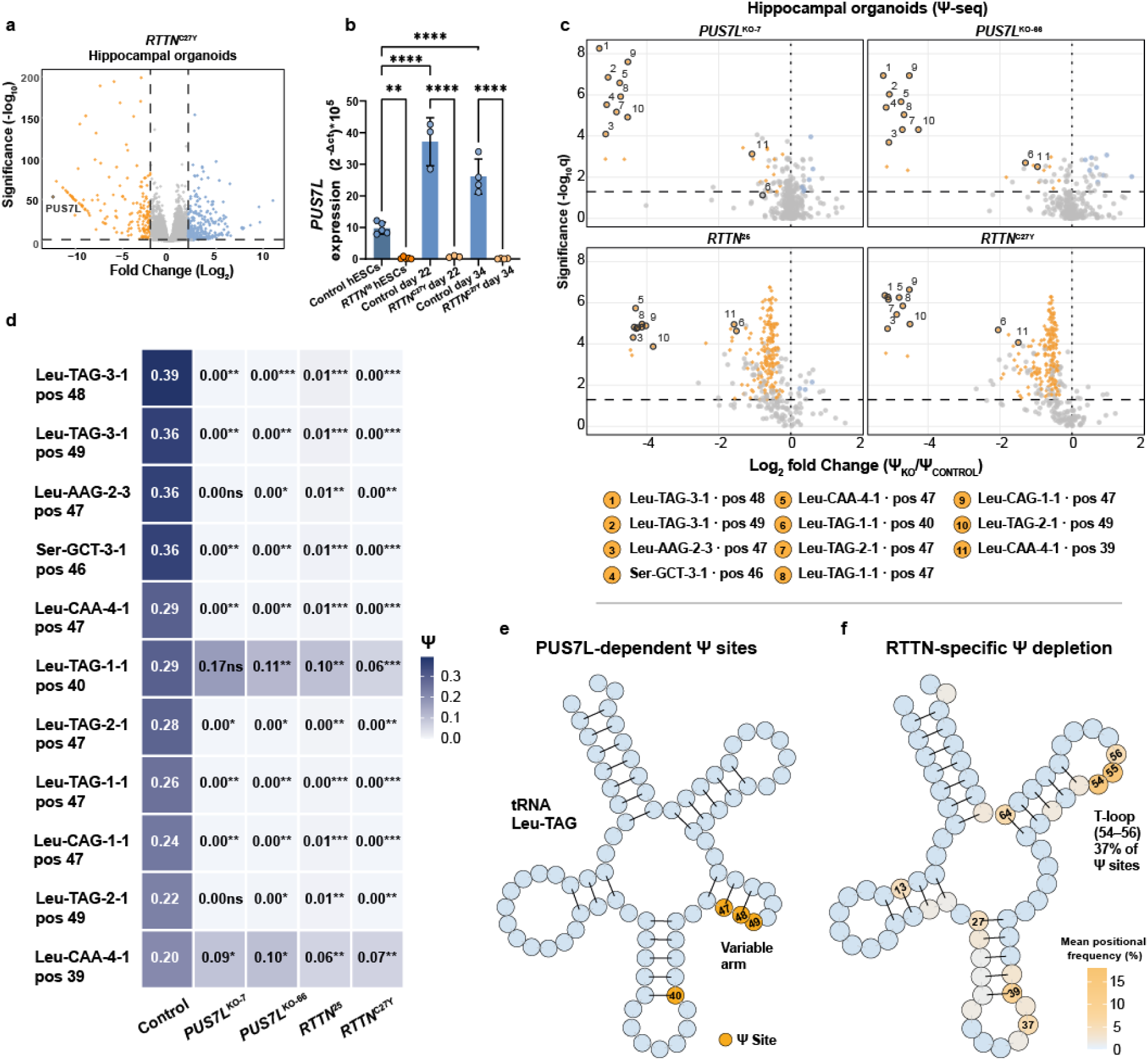
*RTTN* mutations induce distinct PUS7L-dependent and RTTN-specific tRNA pseudouridylation signatures in hippocampal organoids. **a,** Volcano plot of differential gene expression in *RTTN*^C27Y^ hippocampal organoids relative to controls. PUS7L is among the most significantly downregulated genes. **b,** Quantitative RT-PCR validation of *PUS7L* expression in control and *RTTN*^10^-mutant human embryonic stem cells (hESCs) (n=5), *RTTN*^C27Y^ hippocampal organoids day-22 (n=3) and day-34 (n=4) hippocampal organoids. Data are presented as mean ± SD. Statistical significance was determined using one-way ANOVA (F = 70.00, P < 0.0001) followed by Šídák’s multiple comparisons test. P values: P < 0.01 (**), P < 0.0001 (****). **c,** Volcano plots of log₂ fold-change in Ψ (*PUS7L*^KO-7^, *PUS7L*^KO-66^ or *RTTN*^25^ or *RTTN*^C27Y^) versus control -log₁₀(q) for all candidate tRNA uridine (T-reference) positions, one panel per knockout or mutant line, generated using an independent quasi-binomial generalized linear model (GLM) fitted to raw C and T read counts (see Methods). Orange and blue points denote sites significant at q < 0.05 and |ΔΨ| > 0.10 (orange, Ψ loss; blue, Ψ gain), whereas grey points indicate nonsignificant sites. The eleven reproducible PUS7L-dependent sites are outlined and numbered (1-11) and correspond to the sites shown in panel (**d**). The dashed horizontal line indicates q = 0.05. Across the 11-site core signature, BH-corrected q-values ranged from 5.6 × 10⁻⁹ to 0.074 (n=4 for each group). **d,** Heatmap of the mean Ψ level (BACS C/(C+T) conversion fraction) across the 11-site core PUS7L-dependent signature (tRNA-Leu/Ser positions 39/40/46–49), with one row per site and one column per condition. Cell values represent group means (n = 4). Asterisks indicate Benjamini-Hochberg (BH)-corrected q-values from two-sided Welch’s t-tests versus the corresponding control (q < 0.05, q < 0.01, *q < 0.001; ns, not significant). **e,** Structural localization of the shared PUS7L-dependent pseudouridylation sites mapped onto a representative human tRNA-Leu-TAG secondary structure. Highlighted positions correspond to the canonical variable-arm PUS7L target sites (46-49). **f,** Structural distribution of *RTTN*-specific pseudouridylation defects mapped onto tRNA secondary structure. Color intensity represents the mean positional frequency (%) of *RTTN*-specific depleted Ψ sites, calculated as the average percentage of sites identified in *RTTN*^25^ and *RTTN*^C27Y^. *RTTN*-associated Ψ depletion is broadly distributed across the tRNA molecule but is strongly enriched within the T-loop (positions 54-56), which accounts for approximately 37% of all *RTTN*-associated Ψ sites, in contrast to the restricted variable-arm signature observed following PUS7L loss.

The ribosomal-state and biosynthetic defects observed in *RTTN*-mutant organoids (Figs. 2-4) predict downstream consequences for neural progenitor behaviors that depend heavily on protein synthesis, including cell-cycle progression, mitotic timing, and coordinated nuclear and neuronal motility. Consistent with this, *RTTN* expression is enriched in cycling progenitors and varies with cell-cycle state in the developing human brain (Extended Data Fig. 1c), suggesting a tight coupling to proliferative programs *in vivo*. We therefore determined whether *RTTN* mutations link translational dysfunction to defects in progenitor proliferation and tissue dynamics using complementary live-imaging and endpoint assays.

We leveraged brain-on-chip organoids to achieve prolonged live imaging and reliable tracking of individual progenitor divisions. Live imaging revealed a pronounced delay in mitotic progression in *RTTN*^C27Y^ cells. Tracking individual divisions across mitotic phases showed an increase in total mitotic duration and prolongation of specific mitotic intervals compared with control (Fig. 6a). *RTTN*^C27Y^ organoids also exhibited altered interkinetic nuclear migration, with significantly reduced nuclear velocities for basal and apical-directed movements (Fig. 6b). We next extended these findings to hippocampal organoids. Immunostaining demonstrated increased proliferative (Ki67) and mitotic (pHH3) markers, along with elevated apoptotic signaling (cleaved caspase-3) in *RTTN*^C27Y^ organoids (Fig. 6 c,d,e), indicating dysregulated progenitor cell-cycle progression, characterized by accumulation of cycling and mitotic cells, together with increased cell death. Consistent with this, EdU incorporation assays showed a significant reduction in the fraction of S-phase cells across multiple *RTTN* mutant lines (Fig. 6f), supporting reduced proliferative capacity.

**Figure 6.**
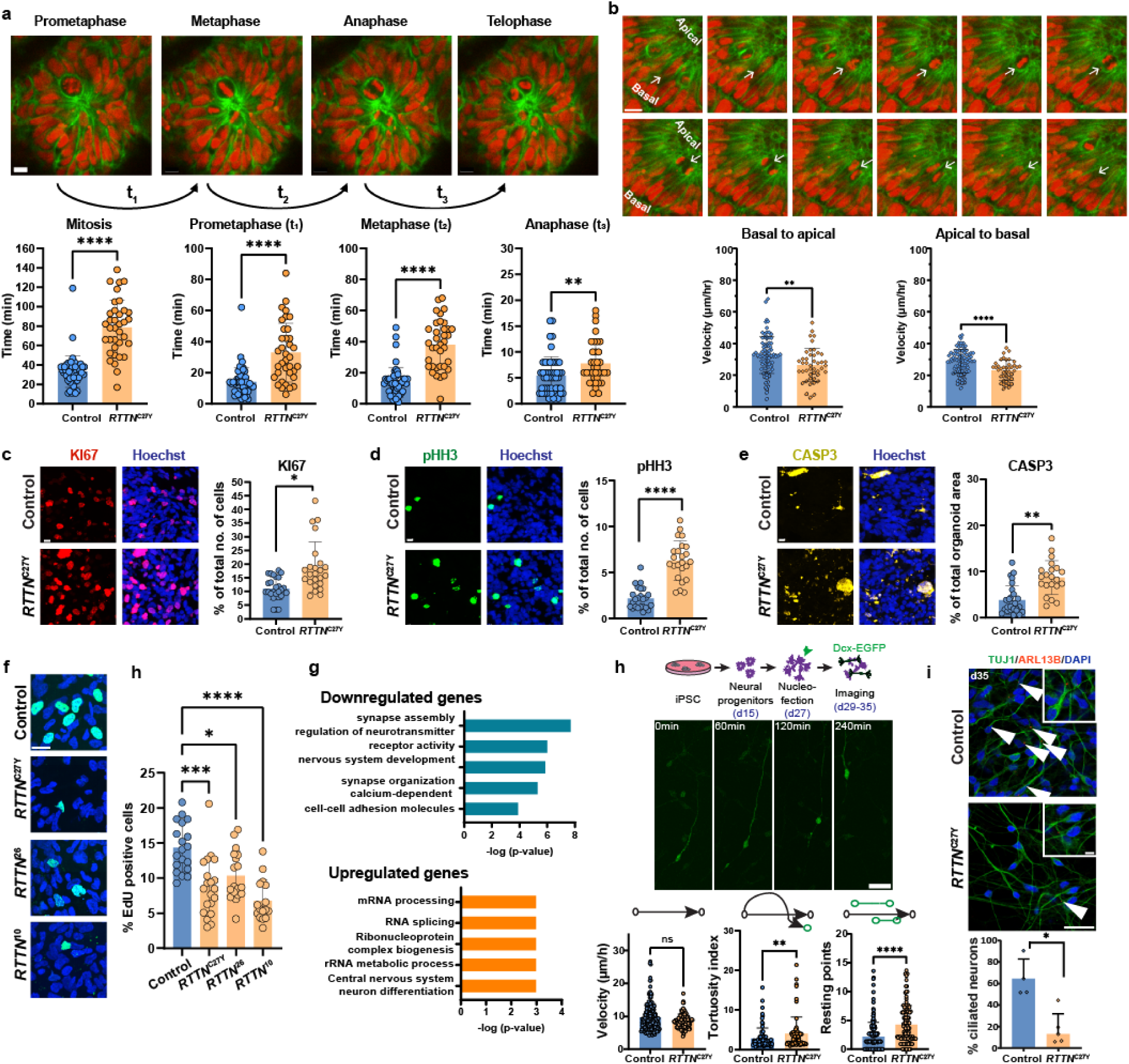
*RTTN* mutations couple reduced biosynthetic programs to progenitor cell-cycle delay, impaired proliferation, and altered neuronal motility. **a,** Representative live-imaging frames showing prometaphase, metaphase, anaphase, and telophase in brain-on-chip organoids (green-Lyn-GFP, red-H2B mCherry). Below, quantification of total mitotic (P<0.0001) duration and phase lengths (prometaphase (t₁) (P<0.0001), metaphase (t₂) (P<0.0001), anaphase (t₃) (P=0.0069) comparing control and *RTTN*^C27Y^. Each dot represents one mitotic cell (Control, n = 50; RTTN^C27Y^, n = 36). Cells were analyzed from organoids at two developmental time points (control days 13–14, mutant days 13-14 and 20–22) and 4-6 movies (batches). Two-tailed t-test, scale bar, 10 µm. **b,** Time-lapse examples of interkinetic nuclear migration (IKNM) in brain-on-chip organoids (green-LYN-GFP, red-H2B mCherry), tracked nuclei indicated. Right, quantification of nuclear velocity for apical-directed (P=0.0033) and basal-directed movements (P<0.0001) in control versus *RTTN*^C27Y^ (Two-tailed Mann–Whitney test), RTTN^C27Y^ (n=44) and control (n=86) cells were analyzed from day 14 and day 20 organoids and 3-4 movies (batches). Scale bar, 15 µm. **c,** Immunostaining in hippocampal organoids for Ki67 (proliferation), P=0.0458, **d,** phospho-histone H3 (pHH3) (mitotic cells), P<0.0001, and **e,** cleaved caspase-3 (apoptosis), P=0.0047, with Hoechst nuclear counterstain. Right, quantification comparing control and *RTTN*^C27Y^. Each dot represents one image. The analysis includes 3-4 sections from 3-4 organoids, from two independent batches. Statistical significance was determined using a nested t-test. Scale bar, 10 µm. **f,** EdU incorporation in hippocampal organoids (day 63, Control, *RTTN*^26^, *RTTN*^C27Y^, *RTTN*^10^). Left, representative images (EdU-positive nuclei shown in green; nuclei in blue). Right, quantification of the percentage of EdU-positive cells across conditions with multiple-comparison statistics as indicated. Each dot represents an individual measurement (Control n = 19; *RTTN*^C27Y^ n = 20; *RTTN*^26^ n = 18; *RTTN*^10^ n = 16). Bars indicate mean ± SEM. Statistical analysis was performed using a Kruskal-Wallis test (P < 0.0001), followed by Dunn’s multiple comparisons test. Adjusted P values: Control vs. *RTTN*^10^, P < 0.0001; Control vs. *RTTN*^26^, P = 0.0301; Control vs. *RTTN*^C27Y^, P = 0.0003. The data shown represents one of two independent experiments with similar results. Scale bar, 20 µm. **g,** Gene Ontology enrichment of differentially expressed genes from hippocampal organoid RNA-seq (comparison and directionality as shown): top (blue) terms correspond to one direction of change and bottom (orange) terms correspond to the opposite direction (e.g., downregulated vs upregulated). **h,** Workflow schematic for the neuronal motility assay (iPSC → neural progenitors → nucleofection → imaging) and representative time-lapse images of a labeled neuron over minutes (0-240 min; fluorescent channel shown). Quantification of neuronal motility features derived from tracking: velocity, tortuosity index, and number of resting points, comparing control and *RTTN*^C27Y^. Each dot represents a tracked cell, bars indicate mean ± SD; significance is indicated for the Mann-Whitney test (tortuosity index, P = 0.009; resting points, P < 0.0001), n=3 independent differentiation batches, except for tortuosity index, where n = 2. Scale bar, 50 µm**. i,** Immunostaining of neuronal cultures at day 35 for TUJ1 (neurons), ARL13B (cilia), and DAPI, comparing control (n=4) and *RTTN*^C27Y^ (n=5), with quantification of the percentage of ciliated TUJ1+ cells (right). Each dot represents the average percentage of ciliated TUJ1+ neurons calculated from a single biological replicate (n = 4 independent batches, Mann-Whitney test P = 0.0159. Scale bar, 25 µm.

At the transcriptional level, RNA-seq analysis revealed pathway-level changes consistent with the cellular phenotypes, including enrichment of neuronal development and morphogenesis terms among downregulated genes and RNA processing-related pathways among upregulated genes (Fig. 6g). In addition to differential gene expression, *RTTN*-mutant organoids exhibited widespread alternative splicing alterations, with enrichment for pathways related to cell junction assembly, neuron development, neurogenesis, and cellular morphogenesis (Extended Data Fig. 5a,b). Notably, a substantial overlap was observed between alternatively spliced genes identified in *RTTN*^C27Y^ and *RTTN*^25^ organoids, consistent with the similar developmental phenotypes observed across the RTTN-mutant lines (Extended Data Fig. 5c). Among the commonly affected targets, *RTTN*-mutant lines displayed altered exon usage at the *ERBB4* locus (Extended Data Fig. 5d,e), a gene implicated in neuronal migration and signaling, highlighting the potential impact of disrupted RNA-processing programs on key neurodevelopmental processes.

Finally, we assessed whether these molecular changes translate to functional deficits using a live-cell motility assay. *RTTN*^C27Y^ neurons displayed altered movement dynamics compared with controls (Fig. 6h), with reduced velocity, increased tortuosity, and more frequent resting periods (Fig. 6h). In parallel, *RTTN*^C27Y^ neurons displayed a reduced fraction of ARL13B-positive cilia on TUJ1-positive neurons (Fig. 6i), consistent with impaired neuronal ciliation. Together, these results link *RTTN*-associated translation ribosome and translational defects to delayed mitotic progression, reduced progenitor proliferation, and altered neuronal motility and ciliation, establishing a mechanistic connection between disrupted ribosome homeostasis and key neurodevelopmental phenotypes.

RTTN has been classically viewed as a centrosome/centriole factor required for proper centrosome maturation and cilia-related signaling, and human *RTTN* mutations are linked to severe malformations of cortical development, including polymicrogyria and microcephaly^2, 3, 5, 6, 12, 13, 19–21^. However, the molecular mechanisms linking primary centrosomal dysfunction to impaired cortical growth have remained poorly understood. Our findings identify an unexpected association among RTTN dysfunction, ribosome homeostasis, tRNA pseudouridylation, and translational control, suggesting that impaired biosynthetic capacity may be a major mechanism underlying neurodevelopmental failure in *RTTN*-associated disease. In addition, *RTTN* mutations induced both a shared *PUS7L*-dependent pseudouridylation signature and a broader *RTTN*-specific defect in tRNA pseudouridylation, further linking *RTTN* dysfunction to disrupted RNA metabolism. Notably, loss of the related pseudouridine synthase *PUS7* also causes microcephaly and intellectual disability in humans^22–24^ and plays a direct role in translation by pseudouridylating tRNA-derived fragments that repress translation initiation in stem cells^25^. This precedent differs mechanistically from our findings: PUS7 acts at distinct sites (including tRNA position 13, 20B, 35, 36, 50 and position 8 at tRNA-derived fragments) and represses translation in trans, so that its loss increases protein synthesis, whereas PUS7L modifies the variable arm of mature tRNAs and the *RTTN*-associated defect instead reduces translational output. PUS7 nonetheless establishes tRNA pseudouridylation as a plausible link between impaired translation and microcephaly. Together, these findings indicate that *RTTN*-associated pathology reflects not only centriole and ciliary defects but also impaired ribosome biogenesis, tRNA pseudouridylation, and translational control, all of which compromise the biosynthetic capacity of neural progenitors during corticogenesis.

A particularly novel finding was the interaction of RTTN with multiple RNA-binding and RNA-processing factors. Beyond serving as microtubule-organizing centers, centrosomes are increasingly recognized as hubs for RNA localization and post-transcriptional regulation. Recent studies have identified spliceosomal proteins, RNA helicases, heterogeneous nuclear ribonucleoproteins (hnRNPs), and exon junction complex (EJC) components at centrosomes and centriolar satellites, where they contribute to centrosome biogenesis, ciliogenesis, and neural stem cell function^26–30^. These observations support a functional link between RTTN and RNA-regulatory processes. The RTTN interactome included several nucleolar proteins involved in ribosome biogenesis, including DDX21, DDX18, FBL, and LYAR, supporting an unexpected association between RTTN and nucleolar pathways regulating rRNA synthesis and processing. Among these, DDX21 is particularly notable because it functions in rRNA biogenesis, dynamically redistributes between nucleolar and nucleoplasmic compartments in response to cellular state, and participates in broader RNA-regulatory processes^31, 32^. Notably, RTTN also interacted with proteins implicated in tRNA maturation, including the tRNA-splicing ligase RTCB, suggesting that RTTN-associated RNA dysregulation extends beyond rRNA metabolism. Moreover, the RTTN-specific T-loop pseudouridylation signature was not accompanied by reduced expression of the canonical T-loop pseudouridine synthases, TRUB1 or PUS10, suggesting that these defects arise from perturbation of the RNA-processing environment or tRNA maturation rather than from direct loss of individual modifying enzymes. At the level of translation dynamics, we observed reduced nascent protein synthesis, altered ribosome-state distributions by polysome profiling, and perturbations in ribosomal subunit balance across multiple *RTTN* mutations and organoid contexts. In hippocampal organoids, the emergence of half-mer-like features, together with shifted subunit ratios, is consistent with defects in ribosome maturation or subunit joining/assembly. Analogous abnormalities have been described in ribosome maturation disorders, including SBDS/EFL1-associated Shwachman-Diamond syndrome^33^, and in mechanistic studies of eIF6 recycling^34^. In telencephalon organoids, transcripts with reduced translational efficiency were enriched for pathways related to cilium assembly, cytoskeletal organization, and neuronal developmental programs, closely matching observed cellular phenotypes. The enrichment of ribosomal protein transcripts within these altered ribosome states is particularly notable, as efficient translation of ribosomal protein mRNAs is required to sustain ribosome biogenesis and translational capacity. Thus, RTTN-associated translational defects may secondarily impair synthesis of core ribosomal components, raising the possibility of a self-reinforcing translational defect in which impaired synthesis of ribosomal components further destabilizes ribosome homeostasis. Altered tRNA pseudouridylation may further contribute to reduced translational efficiency by impairing the structural integrity and ribosomal function of mature tRNAs. Complementary rDNA-spanning RT-qPCR and nascent RNA labeling revealed signatures consistent with disrupted pre-rRNA processing rather than global loss of rRNA transcription, and is consistent with dysregulated or compensatory nucleolar activity in the context of impaired rRNA maturation. Increased 5-EU incorporation further supports elevated nucleolar transcriptional activity, consistent with dysregulated or compensatory nucleolar function in response to impaired rRNA maturation and elevated nucleolar stress. Persistent nucleolar stress is known to engage cell-cycle checkpoint pathways, including nucleolar stress-to-p53 signaling mediated by ribosomal protein-dependent inhibition of MDM2^35, 36^. In rapidly proliferating neuroepithelia, such stress responses may provide a mechanistic connection between impaired ribosome homeostasis and the prolonged mitosis, reduced S-phase entry, and diminished progenitor expansion observed in *RTTN*-mutant organoids.

A second novel aspect of this study is the integration of these translation-associated molecular phenotypes with quantitative developmental readouts. The observed cellular phenotypes, including prolonged mitosis, reduced interkinetic nuclear migration, and altered proliferation and apoptosis markers, are consistent with the well-established vulnerability of neural progenitors to centrosome dysfunction^26, 37, 38^. This vulnerability is further underscored by our finding that *RTTN* is highly expressed in cycling progenitors in the first-trimester human brain, suggesting that *RTTN* function is deployed in precisely the populations most dependent on robust translational capacity. Beyond its established centrosomal role, our findings identify ribosome homeostasis and RNA metabolism as major processes disrupted by *RTTN* mutations. Transcriptome analyses further reveal altered RNA-processing programs, including alternative splicing of neurodevelopmental genes such as *ERBB4*. Together with emerging evidence that centrosomes coordinate multiple aspects of post-transcriptional regulation^26–30^, these findings suggest that *RTTN* contributes to RNA metabolism at several levels during human corticogenesis.

Overall, these findings point to broad disruption of ribosome-state regulation and translational engagement in *RTTN*-mutant organoids, although the molecular mechanisms linking centrosome-associated RTTN to nucleolar and translational regulation remain to be defined. It will be important to determine whether these effects arise through altered cell-cycle progression, nucleolar organization, stress signaling, or more direct coupling between centrosome state and ribosome biogenesis, and whether restoring ribosome homeostasis can ameliorate progenitor and neuronal phenotypes in *RTTN*-mutant models. Together, our findings support an emerging view of the centrosome as an organizer of post-transcriptional gene regulation, extending its functions beyond microtubule organization to include ribosome biogenesis, tRNA pseudouridylation, translational control, and RNA processing during human corticogenesis. Our model suggests testable points of convergence, including ribosome-state regulation, nucleolar stress signaling, and selective translational dysregulation, that may represent actionable mechanisms for future therapeutic exploration in *RTTN*-related neurodevelopmental disorders.

## Acknowledgments

We would like to thank Isabella Zagorski for her help in karyotyping the iPSC lines, and Andrea Steiner-Mezzadri, Sabine Ulbricht, and Martina Buerkle for their help with cell culture and immunoprecipitation. We are especially grateful to Prof. Grazia Mancini for providing patient fibroblast cell lines. We would like to sincerely thank Tamar Sapir and Anna Gorelik for their valuable assistance with cell culture experiments and insightful scientific discussions.

