## supplementary figures and methods for "RTTN moonlights beyond the centrosome to control ribosome biogenesis and tRNA modification in human brain organoids"

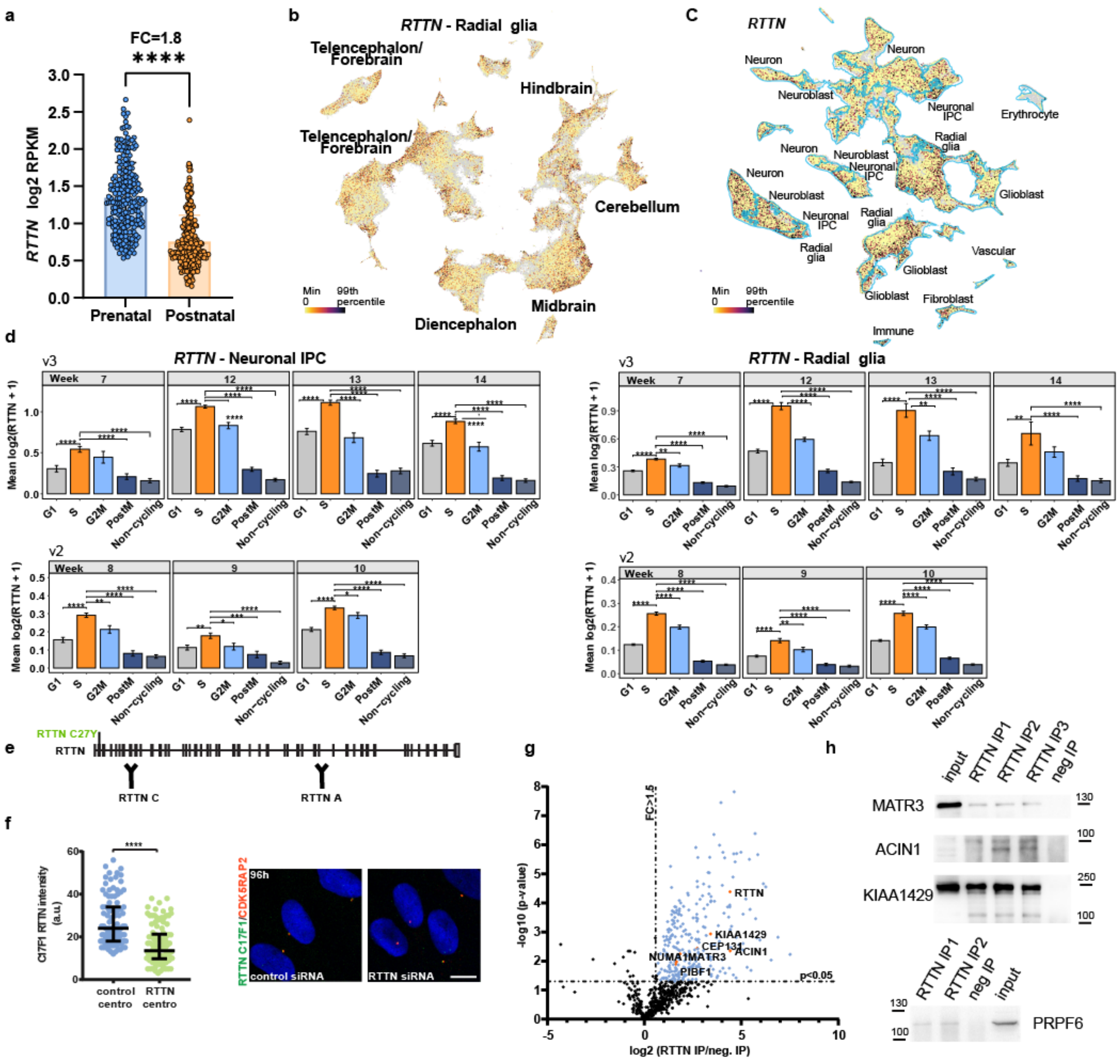

**Extended Data Fig. 1: *RTTN* expression, cellular distribution, and molecular interactions during human corticogenesis.**

**a,** *RTTN* mRNA expression across human postmortem brain development from 7-37 post-conception weeks (PCW) compared with postnatal ages (3 months to 40 years; prenatal *n* = 237; postnatal *n* = 287), from the BrainSpan Atlas of the Developing Human Brain. Prenatal samples show significantly higher *RTTN* expression than postnatal samples (two-sided t test, *P* <0.0001). **b,** UMAP projection of single-cell transcriptomes from human fetal brain showing *RTTN* expression across major regions (telencephalon/forebrain, diencephalon, midbrain, hindbrain, and cerebellum), with enrichment in radial glia populations. **c,** UMAP projection of single-cell RNA-seq from first-trimester developing human brain (PCW 5, 6, 7, and 11; Braun et al., Science 2023[1]) showing *RTTN* expression overlaid on major annotated cell populations (e.g.,

radial glia, neuronal IPCs, neuroblasts/neurons, and non-neuronal populations).
**d,** Mean *RTTN* expression (log2(*RTTN* + 1)) in neuronal IPCs and radial glia stratified by cell-cycle phase and developmental week. Analyses were performed separately for Chromium v2 and v3 datasets. Bars represent mean expression, and error bars indicate SEM. Statistical comparisons were performed between S-phase cells and the remaining cell-cycle phases using two-sided Wilcoxon rank-sum tests with Benjamini-Hochberg correction for multiple testing. Expression is highest in S and G2/M phases. **e,** Schematic of RTTN protein domain organization indicating the epitopes targeted by RTTN antibodies. **f,** ICC antibody validation. Quantification of RTTN signal intensity at centrosomes in control and RTTN-depleted fibroblasts. siRNA-mediated *RTTN* knockdown reduces RTTN levels at the centrosome ($P < 0.0001$, Mann-Whitney test from 3 independent transfections), Scale bar, 10 μm. **g,** Volcano plot of proteins identified by RTTN immunoprecipitation followed by mass spectrometry. Significantly enriched interactors (blue; $P < 0.05$, fold change > 1.5) include centrosomal and RNA-associated proteins (for example, CEP131, KIAA1429, ACIN1, MATR3, NUMA1, and PIBF1). **h,** Immunoblot validation of RTTN-interacting proteins following immunoprecipitation, confirming RTTN association with MATR3, ACIN1, KIAA1429, and PRPF6 compared to control immunoprecipitation in NPCs-derived lysates. IPs performed on lysates from 3 independent batches of NPC differentiations, except for PRPF6 (n = 2).

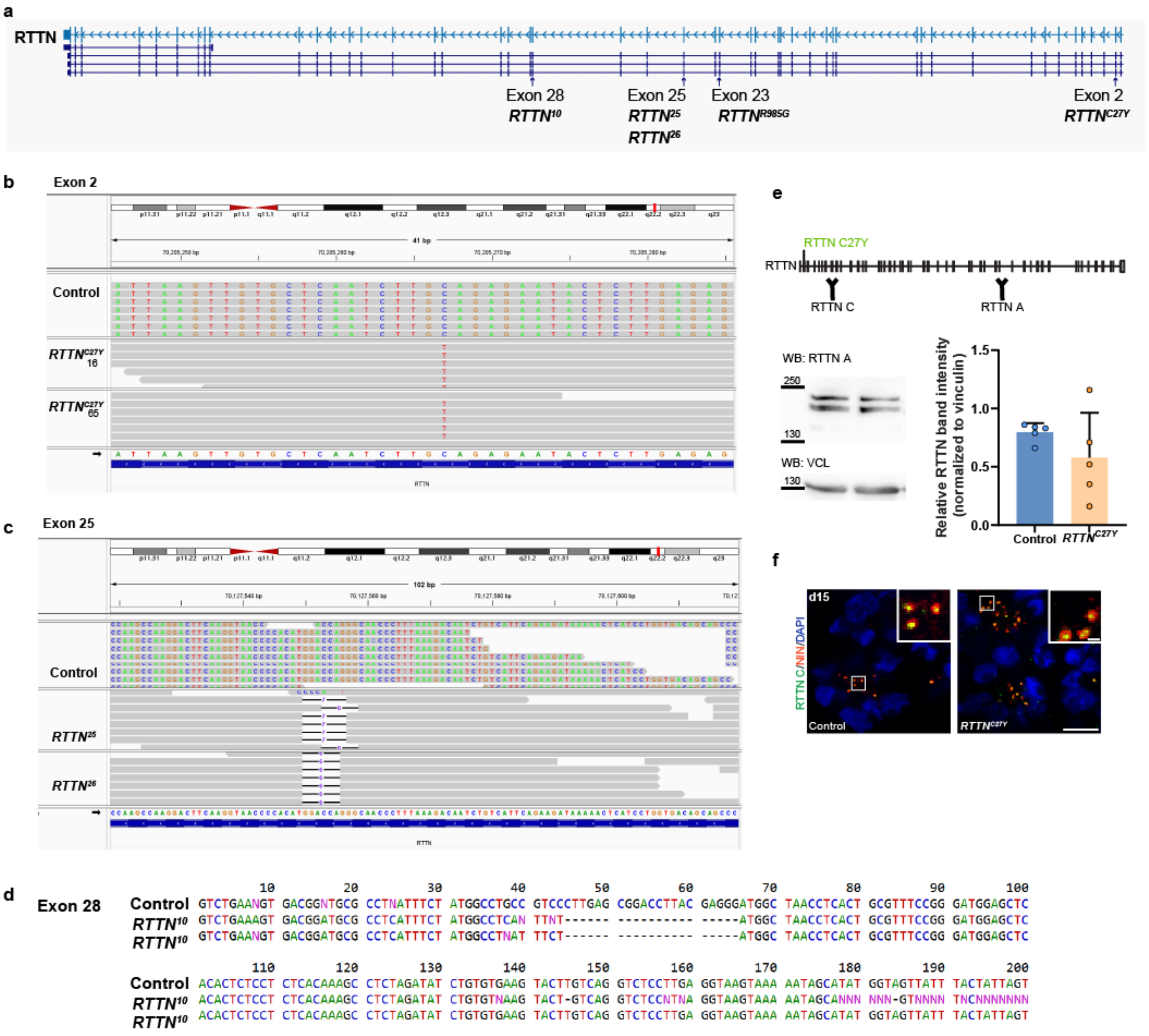


**Extended Data Fig. 2: CRISPR-Cas9-mediated editing of *RTTN* and validation of mutant lines.** CRISPR-Cas9 genome editing was used to introduce targeted mutations in *RTTN,* generating mutant clones for downstream functional analyses. Edited lines were validated by RNA-sequencing and Sanger sequencing, confirming the presence of the intended nucleotide substitutions and frameshift-inducing deletions relative to control cells. **a,** Schematic representation of the human *RTTN* gene showing the locations of the patient-derived iPSC mutations *RTTN*$^{R985G}$ (Exon 23) and *RTTN*$^{C27Y}$ (Exon 2), as well as the CRISPR/Cas9-engineered mutant alleles *RTTN*$^{10}$ (Exon 28), *RTTN*$^{25}$, and *RTTN*$^{26}$ (Exon 25) generated in human embryonic stem cells (hESCs). In addition, two independent CRISPR/Cas9-engineered *RTTN*$^{C27Y}$ hESC lines (16 and 65) were generated to model the patient-associated C27Y variant. **b,** IGV browser view of RNA-sequencing reads at *RTTN* exon 2 showing introduction of the Cys27Tyr

(*RTTN*$^{C27Y}$) mutation. The numbers below (16 or 65) represent individual clones. Sequencing reads from the control, and edited lines are aligned to the reference genome. Edited lines show the expected nucleotide substitution, whereas control cells retain the wild-type sequence. **c,** IGV browser view of RNA-sequencing reads validating CRISPR-Cas9 editing at *RTTN* exon 25. Aligned reads from control and independent edited clones (*RTTN*$^{25}$ and *RTTN*$^{26}$) show deletion mutations at the target site. **d,** Sanger sequencing of *RTTN* exon 28 in control and edited lines (*RTTN*$^{10}$ clones), confirming the presence of frameshift-inducing deletions relative to the wild-type sequence. **e,** Schematic of the RTTN protein showing the location of the *RTTN*$^{C27Y}$ variant and the epitopes recognized by the RTTN-C and RTTN-A antibodies. Representative western blot and quantification of RTTN protein levels normalized to vinculin (VCL) in control and *RTTN*$^{C27Y}$ d15 NPC samples. Data from 5 independent differentiation batches. **f,** Representative immunofluorescence images of day 15 NPCs stained for RTTN (green) and NIN (red), with DAPI (blue). Insert show magnified views of the boxed regions. Scale bar 10 µm, inset 1 µm.

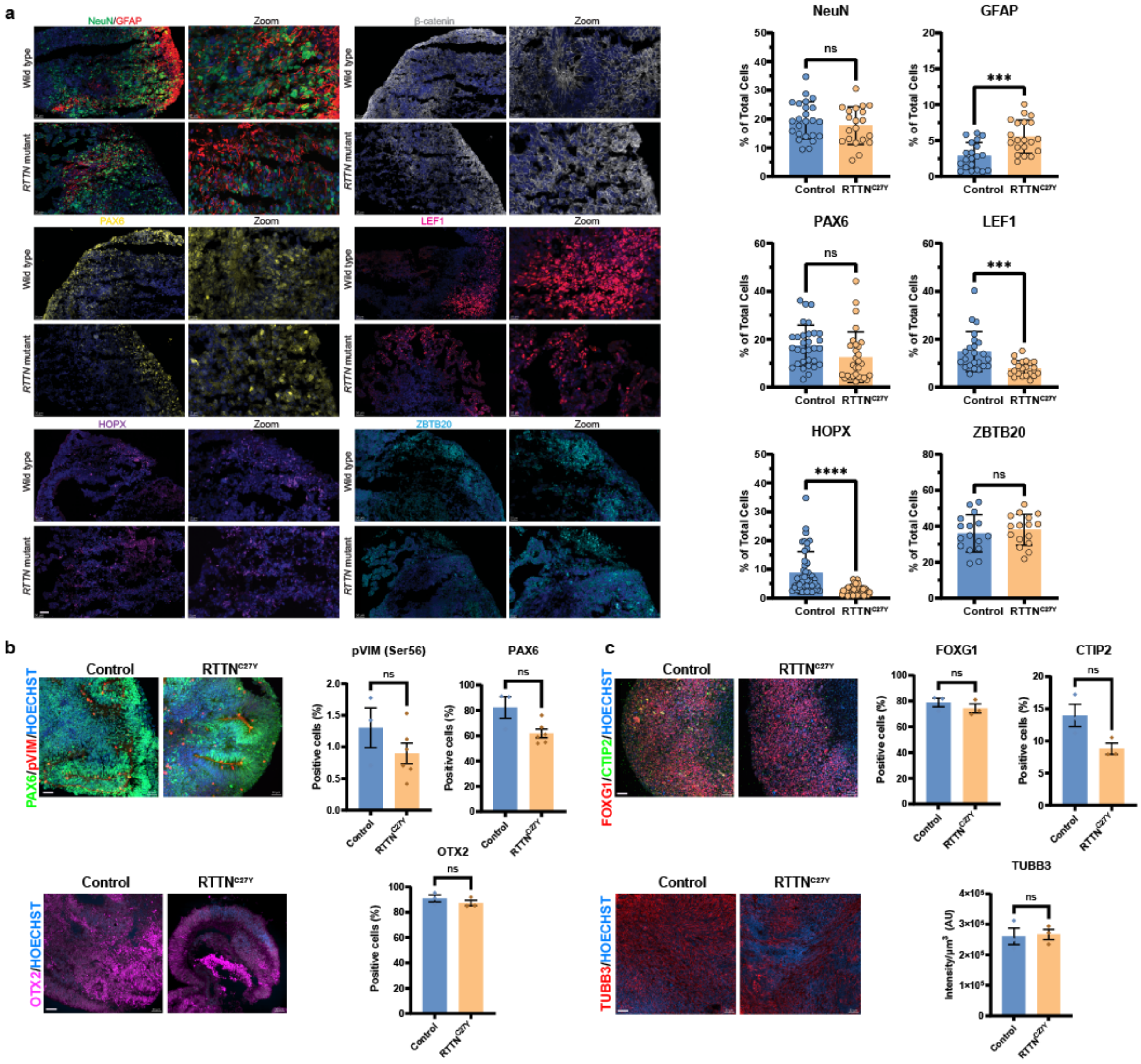

**Extended Data Fig. 3:** Characterization of hippocampal and telencephalic organoids. **a,** Representative immunofluorescence images of day 64 hippocampal organoids showing the expression of hippocampal and neuronal markers. Quantification of marker expression is shown adjacent to the representative images. Each dot represents one image. The analysis includes 3-4 sections from 3-4 organoids, from two independent batches, and Statistical analysis was performed using an unpaired two-tailed t-test with Welch's correction. **b-c**, Characterization of human telencephalic organoids. b, Representative immunofluorescence images of day 30 organoids showing robust neuroepithelial organization and expression of the neural progenitor markers PAX6, pVIM, and OTX2. (Control, n = 3 organoids; *RTTN*$^{C27Y}$, n = 6 organoids.) **c**, Representative immunofluorescence images of day 60 organoids showing telencephalic progenitors marked by FOXG1, cortical neurons marked by CTIP2, and neuronal processes stained with TUBB3. Each dot represents the average from one organoid (Control, n = 3 organoids; RTTN$^{C27Y}$, n = 3 organoids). For each condition, three independent organoids were analyzed. Statistical analysis was performed using an unpaired two-tailed t-test with Welch's correction. Scale bar, 50 μm.

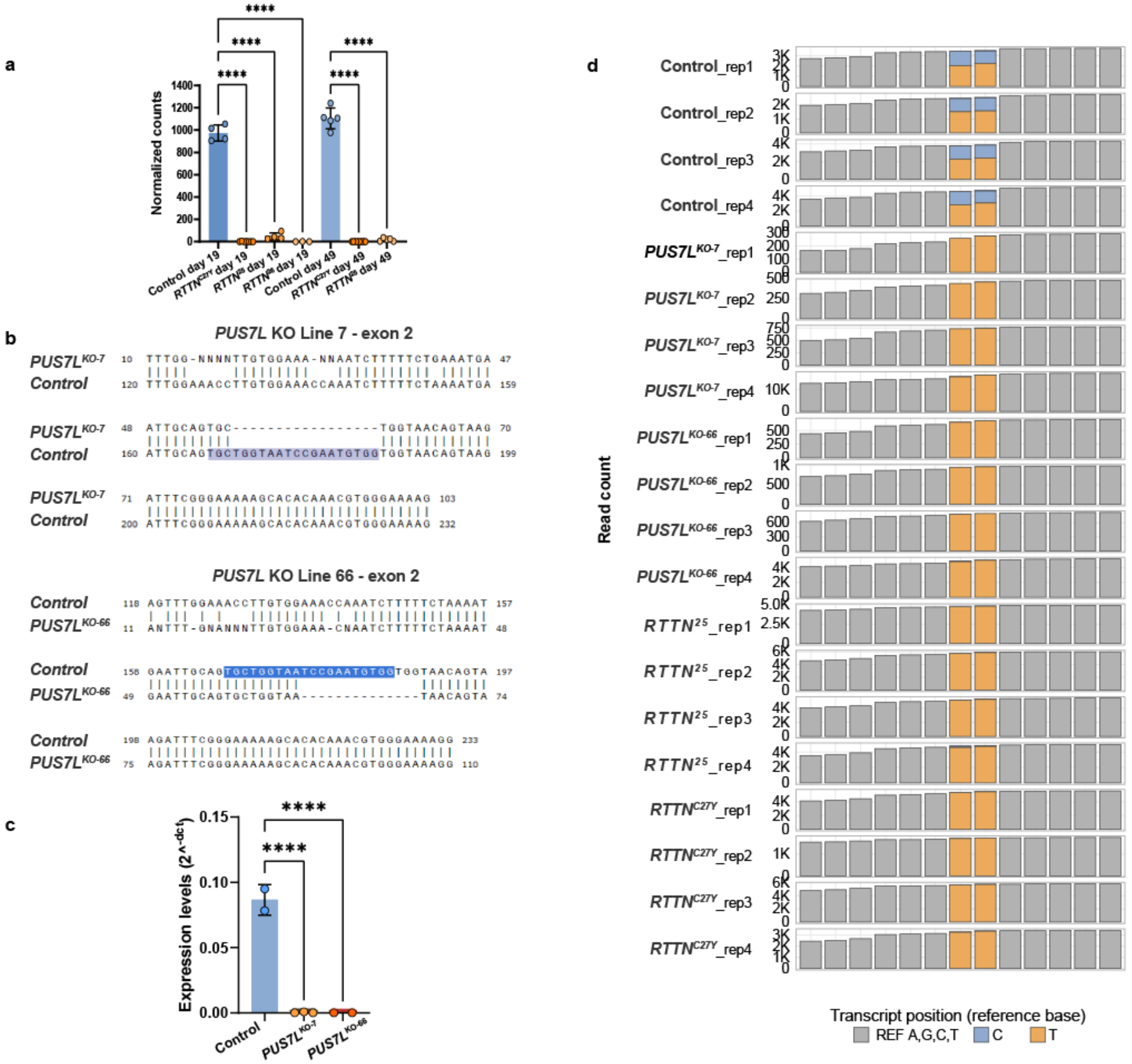


**Extended Data Fig. 4**. Generation and validation of *PUS7L* knockout hESC lines and visualization of BACS-derived Ψ signals. **a,** Normalized RNA-seq counts showing significantly reduced *PUS7L* expression in *RTTN*$^{C27Y}$, *RTTN*$^{25}$ and *RTTN*$^{26}$ hippocampal organoids compared with control organoids. Bars represent mean ± SD (n = 4 (Control day 19), n = 7 (*RTTN*$^{C27Y}$ day 19), n = 4 (*RTTN*$^{25}$ day 19), n = 3 (*RTTN*$^{26}$ day 19), n = 5 (Control day 49), n = 10 (*RTTN*$^{C27Y}$ day 49), and n = 5 (*RTTN*$^{25}$ day 49).). Statistical significance was determined using one-way ANOVA with Šídák's multiple-comparison test (**, P < 0.01; ****, P < 0.0001). **b,** Sanger sequencing of CRISPR/Cas9-edited

*PUS7L* alleles in the two independent knockout lines showing frameshift-inducing insertions/deletions within exon 2 compared with the wild-type control sequence. The CRISPR target region is highlighted. **c,** RT-qPCR validation of *PUS7L* expression in undifferentiated hESCs, confirming efficient knockout in both *PUS7L*$^{KO\text{-}7}$ and *PUS7L*$^{KO\text{-}66}$ lines. Statistical analysis was performed using ordinary one-way ANOVA followed by Dunnett's multiple comparisons test (Control as the reference group). Sample sizes were n = 2 (Control), n = 3 (*PUS7L*$^{KO\text{-}7}$ and n = 2 (*PUS7L*$^{KO\text{-}66}$). All comparisons versus Control were significant (****P < 0.0001). **d,** Hippocampal organoids per-position nucleotide-identity read coverage across Homo sapiens tRNA-Leu-TAG-3-1 (transcript positions 42–54) for all 20 BACS Ψ-seq libraries (Control, *PUS7L*$^{KO\text{-}7}$, *PUS7L*$^{KO\text{-}66}$, *RTTN*$^{KO25,}$ and *RTTN*$^{C27Y}$; n = 4 biological replicates per condition). Grey bars indicate total read depth at reference-matching positions. At the two *PUS7L*-dependent Ψ sites (positions 48 and 49), bars are partitioned into unconverted thymidine (T, yellow) and BACS-converted cytidine (C, blue) reads, with the C/(C+T) fraction representing the pseudouridine (Ψ) signal at each position in each biological replicate.

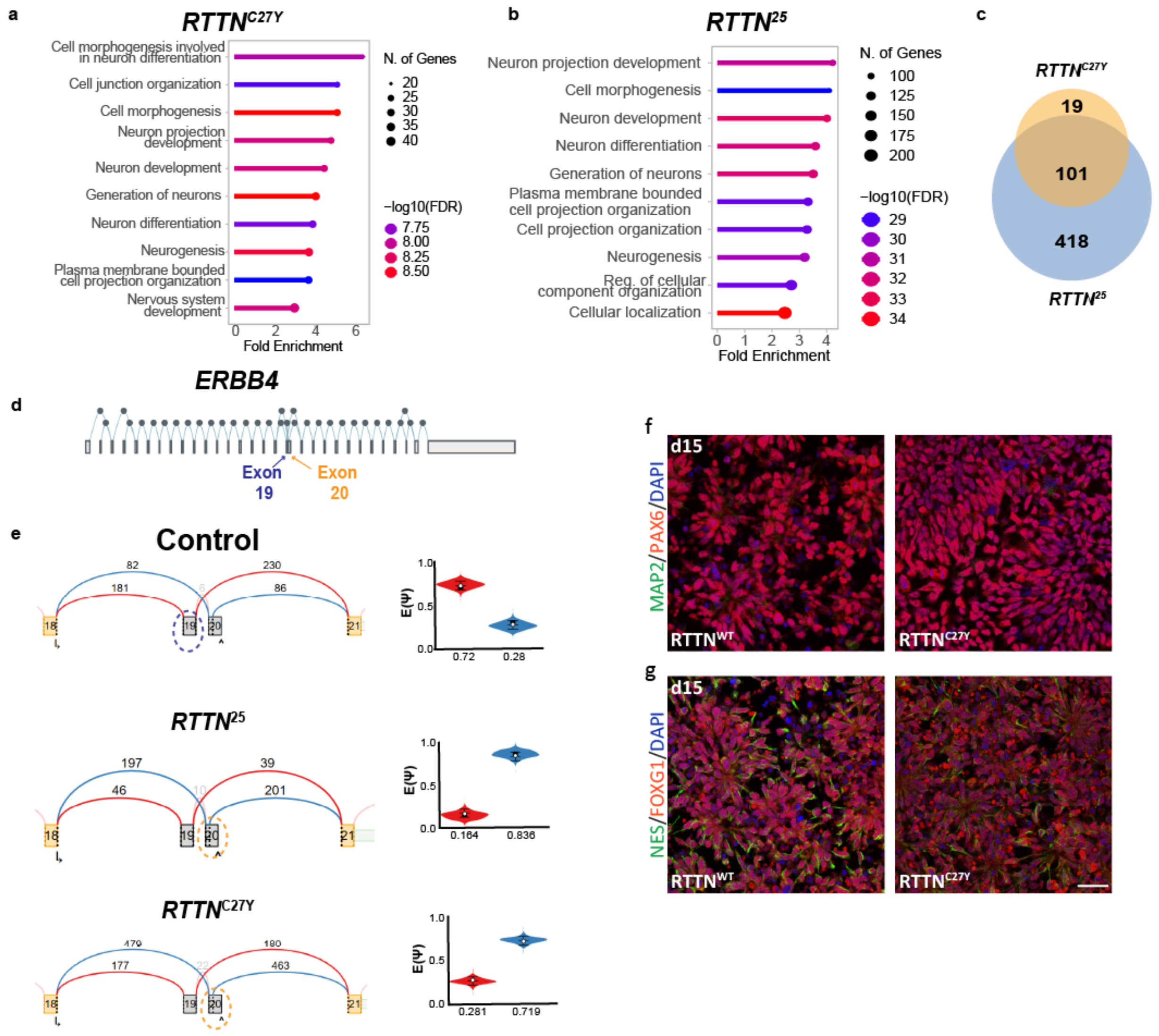


**Extended Data Fig. 5: Alternative splicing alterations in *RTTN* mutant cells converge on neurodevelopmental pathways. a,b** Gene ontology (GO) enrichment analysis of differentially spliced genes in *RTTN*$^{C27Y}$ and *RTTN*$^{25}$ organoids. Dot size indicates the number of genes associated with each GO term, and color denotes enrichment significance (−log10(FDR)). **C,** Overlap of differentially spliced genes between *RTTN*$^{C27Y}$ and RTTN$^{25}$ organoids. **d,** Schematic of the *ERBB4* locus highlighting alternatively spliced exons (exons 19 and 20). **e,** Plots illustrating alternative splicing events in *ERBB4* across control, *RTTN*$^{25}$, and *RTTN*$^{C27Y}$ samples. Arcs indicate exon-exon junction reads, and numbers denote read counts. Differential exon usage highlights altered exon inclusion patterns in *RTTN* mutant lines, with corresponding exon inclusion levels (Ψ) shown. Differential splicing events were identified using MAJIQ with thresholds of P(ΔΨ > 0.15) > 0.95. **f,g** Immunofluorescence analysis of neural differentiation at day 15 (d15). **f,** MAP2 (green), PAX6 (red), and DAPI (blue). **g,** NES (green), FOXG1 (red), and DAPI (blue). Representative images from control and RTTN p.Cys27Tyr (C27Y) cells show comparable expression of neural progenitor and neuronal markers. Scale bars, 50 µm.

### Materials and Methods

#### Ethics statement

Work with hESCs (WIBR3, NIHhESC-10-0079) and iPSCs was carried out with approval from the Weizmann Institute of Science IRB (Institutional Review Board), and genome editing was performed. Animal experiments were conducted in accordance with the German animal welfare law and performed with permission and in accordance with all relevant guidelines and regulations of the district government of Upper Bavaria (Bavaria, Germany; Animal protocol number ROB-55.2Vet-2532.Vet_03-17-68).

#### Human fibroblast culture

Human fibroblasts were maintained in DMEM/GlutaMAXTM (Gibco) supplemented with 10% qualified Fetal Bovine Serum (Gibco 10500056), 0.01M HEPES, 1% NEAA, and 1% Penicillin/Streptomycin (both Life Technologies). For knockdown experiments, cells were seeded at ($5{,}3*10^4$ cells/cm$^2$) and transfected with ON-TARGETplus SMART pool siRNAs against RTTN (Dharmacon, L-031139-01-0005) or non-targeting control siRNA (Dharmacon, D-001810-10-05) using the Dharmafect3 reagent (Dharmacon, T-2003-01) according to the manufacturer's instructions. Transfected cells were collected 96h after transfection for Western blot analysis or fixed with 100% MeOH at -20°C for 10min.

#### Generation and maintenance of human iPSCs

Induced pluripotent stem cell (iPSC) lines from control or from a patient carrying a homozygous RTTNC27Y and pArg985Gly mutations were reprogrammed from fibroblasts at the iPSC Core Facility, Helmholtz Center Munich. Fibroblasts were either purchased (CRL-2522, ATCC) or obtained with informed patient consent from the Department of Clinical Genetics at Erasmus MC, the Netherlands. Isolated iPSC clones were verified for pluripotency and normal karyotype and maintained on Geltrex-coated plates (Life Technologies A1413302) in mTeSR1TM medium (StemCell Technologies) at 37°C, 5% CO2, and ambient oxygen levels with daily medium changes. When cells reached 90% confluency, they were shortly incubated with Collagenase Type IV

(StemCell Technologies, 5-7min), scratched to dissociate colonies, and then distributed to new plates at a 1:6-1:12 ratio.

**Human embryonic stem cells**

Human embryonic stem cells (hESCs), NIH line NIHhESC-10-0079 (WIBR3) were maintained on mouse embryonic fibroblast (MEF) or on Matrigel (Corning 354234) -coated tissue culture plates in human naïve medium[2] supplemented with 10 µM ROCK inhibitor (Medchem express HY-10583) for 24 hours at 37°C with 5% CO2. Cells were passaged every 4–5 days using Trypsin-EDTA (0.05%) or TrypLE™ Express Enzyme (Thermo Fisher Scientific 12604013) according to the manufacturer's instructions. All cell lines were routinely tested for mycoplasma contamination and maintained under standard pluripotent stem cell culture conditions.

**Dorsal forebrain cell generation and culture**

Neural cells were generated as previously published[3] with some modifications. Briefly, iPSCs were dissociated with StemPro Accutase Cell Dissociation Reagent (Gibco A1110501) to single cells, and 140000-280000 cells/cm2 were plated on Matrigel-coated (Corning 354230) dishes in mTeSR1TM medium supplemented with 10 µM Rho-associated kinase inhibitor Y-27632(2HCl) (ROCK, StemCell Technologies 72304) (day 0 of differentiation, d 0). The next day, when cells were fully confluent, neural induction was initiated by changing the culture medium to N3 medium (50:50 DMEM/F12: Neurobasal, 1% GlutaMAX, 1% B27 with vitamin A, 0.5% N2, 50 µM 2-mercaptoethanol, 0.5% MEM-NEAA, 1% penicillin/streptomycin (all Gibco), and 2.5 µg/ml insulin (Sigma-Aldrich I9278). For the first 10 days, the medium was supplemented with 1µM Dorsomorphin (Sigma-Aldrich P5499) and 10µM SB431542 (Sigma-Aldrich S4317) and replaced daily. On d10 cells were passaged with StemPro Accutase Cell Dissociation Reagent as small clusters and replated in N3 medium with 10 µM ROCK onto poly-ornithine and laminin-coated plates (Sigma-Aldrich L2020) at a 1:2-1:4 ratio. Media was changed to N3 without small molecules for the next two days, and then, on d14, before splitting on d15, to N3 with ROCK only. Media was then replaced with fresh N3 without ROCK on d16, and cells were passaged every 5-6 days as above when they reached confluence (e.g., d21, d27). After d29, the cells were not split to induce terminal differentiation.

**CRISPR-Cas9 genome editing and generation of *RTTN* mutant lines**

CRISPR-Cas9 genome editing was used to generate multiple *RTTN* mutant cell lines. Human embryonic stem cells were dissociated into single cells and electroporated with plasmids encoding Cas9, guide RNAs, and a puromycin selection plasmid to enrich for transfected cells.

The $RTTN^{C27Y}$ mutant line carries the patient-associated GRCh38.p13 chr18:70205267 C>T substitution in exon 2 of *RTTN*, resulting in a p.Cys27Tyr missense mutation associated with polymicrogyria (PMG). This mutation was originally identified in a patient with polymicrogyria, seizures, and severe intellectual disability, and affects a highly conserved residue within the first Armadillo domain of RTTN. A guide RNA targeting exon 2 (5′-GAGAGCGCGCTCCCTGATCT-3′) was cloned into the pX330-U6-Chimeric_BB-CBh-hSpCas9 vector (Addgene plasmid #42230) expressing SpCas9 and the chimeric sgRNA under the U6 promoter, and a single-stranded oligodeoxynucleotide (ssODN) donor template containing the pathogenic mutation was co-transfected to enable homology-directed repair.

Additional *RTTN* mutant lines targeting exons 25 and 28 were generated using CRISPR-Cas9-mediated indels to introduce loss-of-function frameshift mutations. These regions were selected based on previously reported *RTTN* patient mutations associated with severe neurodevelopmental phenotypes, including primary microcephaly and intellectual disability. The following guide RNAs were used: exon 25, 5′-AGGTAACCCCACATGGACCA-3′; exon 28, 5′-CCCTTGAGCGGACCTTACGA-3′.

Individual colonies were manually picked, expanded, and screened for successful genome editing by PCR amplification and Sanger sequencing of the targeted *RTTN* loci. Editing outcomes were further validated by RNA sequencing and analysis in Integrative Genomics Viewer (IGV, Broad Institute). Independent *RTTN*-mutant clones, including $RTTN^{C27Y\text{-}4}$, $RTTN^{C27Y\text{-}20}$, $RTTN^{C27Y\text{-}16}$, $RTTN^{C27Y\text{-}65}$, $RTTN^{25}$, $RTTN^{26}$, and $RTTN^{10}$ lines, were used for downstream analyses.

### Generation of *RTTN*-HaloTag knock-in hESCs

To generate endogenous HaloTag-labeled *RTTN* cell lines, CRISPR/Cas9-mediated homology-directed repair (HDR) was performed in hESCs. *RTTN* pCys27Tyr mutant lines (clones 16, 20, and 65) and the control line WIBR3 were used.

A single guide RNA (sgRNA) targeting the *RTTN* locus on chromosome 18 was designed and cloned into the pX330-U6-Chimeric_BB-CBh-hSpCas9 vector (Addgene plasmid #42230), expressing hSpCas9 and the chimeric guide RNA under the U6 promoter. The sgRNA sequence was inserted into the BbsI restriction site according to the Zhang laboratory cloning strategy.

A donor plasmid containing the HaloTag® (Promega) Coding Sequence flanked by *RTTN* left and right homology arms was generated. The HaloTag sequence was fused to *RTTN* through a flexible Gly-Ser-Gly-Gly linker to preserve protein folding and function. The donor construct also contained the corresponding *RTTN* genomic homology regions to facilitate precise insertion at the endogenous locus.

hESCs were dissociated into single-cell suspensions using Accutase and electroporated with the Cas9/sgRNA plasmid and the RTTN-HaloTag donor. Following transfection, cells were plated onto MEF plates in Naive medium supplemented with ROCK inhibitor (Y-27632) (Medchem Express HY-10583) for 24 h to enhance survival.

After recovery and expansion, individual colonies were manually picked and screened for correct HaloTag integration by PCR amplification across the insertion site, followed by Sanger sequencing. Positive clones were further validated for maintenance of pluripotency morphology and HaloTag expression.

### Validation of *RTTN*-HaloTag Knock-In Cell Lines

Correct integration and functionality of the RTTN-HaloTag fusion protein were validated by fluorescence-activated cell sorting (FACS), live-cell HaloTag ligand labeling, and fluorescence microscopy.

Following CRISPR/Cas9-mediated knock-in, cells were incubated with Janelia Fluor® 646 HaloTag® Ligand (50 nM) for 18 h at 37°C. Labeled cells were dissociated into single cells and subjected to fluorescence-activated cell sorting to enrich for HaloTag-positive populations. Sorted cells were expanded and screened for stable HaloTag expression.

For localization studies, cells were transfected with pEGFP-CETN2 (plasmid #2415) to visualize centrioles and centrosomes. Fluorescence imaging demonstrated a discrete signal from the Janelia Fluor® 646 HaloTag® Ligand (Promega GA1121) localized to centrosomal structures and co-localizing with CETN2-positive puncta, consistent with endogenous RTTN localization. Nuclear DNA was counterstained with Hoechst 33342 (Thermo Fisher Scientific H3570).

### Generation of PUS7L-knockout WIBR3 human embryonic stem cells

Genome editing of the PUS7L gene was performed in the WIBR3 human embryonic stem cell (hESC) line using the CRISPR/Cas9 system. Two single-guide RNAs (sgRNAs) targeting coding regions of PUS7L were designed to target exon 2 and exon 3. The target sequences were 5′-TGCTGGTAATCCGAATGTGG-3′ (exon 2) and 5′-CCTGCATAACTAAAATCCGA-3′ (exon 3). Complementary oligonucleotides containing BbsI-compatible overhangs were annealed and cloned into the BbsI site of the pX330-U6-Chimeric_BB-CBh-hSpCas9 vector (Addgene plasmid #42230), which co-expresses SpCas9 and the sgRNA. Cells were transfected with the CRISPR/Cas9 plasmids using electroporation according to the manufacturer's instructions. Following transfection, cells were allowed to recover and plated at low density to permit isolation of single-cell-derived colonies. Individual colonies were manually picked and expanded for molecular characterization.

Genomic DNA was extracted from individual clones, and the genomic regions flanking the CRISPR target sites were amplified by PCR using locus-specific primers. PCR products were analyzed by Sanger sequencing to identify insertions or deletions (indels) generated by non-homologous end joining and to confirm successful genome editing. In addition, total RNA was isolated from edited clones, reverse-transcribed into cDNA, and PUS7L transcript levels were quantified by quantitative real-time PCR (RT-qPCR) using gene-specific primers. Clones carrying biallelic frameshift mutations and exhibiting markedly reduced PUS7L mRNA expression were selected for subsequent experiments. Primer sequences were PUS7L-F 5'-GGTAGACGGGGTCGAGTTTC; PUS7L-R 5'-CCTCAATGCCAACCTCGTCA.

**Generation of monoclonal antibodies against human RTTN**

Lou/c rats were immunized both subcutaneously and intraperitoneally with 40 µg of ovalbumin-coupled peptides of human Rotatin (Peps4LS, Heidelberg), RTNA (aa 1532-1544; APSRTSQDRDPSS), and RTNC (aa 318-330; RTGQRPRGDGQDW), together with 5 nmol CpG (TIB MOLBIOL, Berlin, Germany) and an equal volume of Incomplete Freund's Adjuvant (IFA; Sigma, St. Louis, USA). A boost injection was given 12 weeks later without Freund's adjuvant, and spleen cells were fused with P3X63Ag8.653 myeloma cells using polyethylene glycol 1500 according to standard procedure[4]. After fusion, the cells were plated in 96-well plates using RPMI 1640 (Sigma-Aldrich) with 20% fetal calf serum (FBS, Capricorn Scientific), pyruvate, non-essential amino acids (Pyruvate, NEAA; Sigma-Aldrich), and HAT media supplement (Hybri-Max, Sigma-Aldrich). Hybridoma supernatants were screened in a solid-phase enzyme-linked immunosorbent assay (ELISA) for binding to his-tagged biotinylated peptides coated to streptavidin plates. After blocking with 2% FCS in PBS, hybridoma supernatants were added for 30 min. After one wash with PBS, bound antibodies were detected with a cocktail of HRP-conjugated monoclonal secondary antibodies against the four rat IgG isotypes (TIB173 IgG2a, TIB174 IgG2b, TIB170 IgG1, all from ATCC, R-2c IgG2c homemade). HRP was visualized with ready-to-use TMB substrate (1-StepTM Ultra TMB-ELISA, Thermo). Positive supernatants were further validated by Western blot (WB), immunofluorescence (IF), and immunoprecipitation (IP). Hybridoma cells from selected supernatants were subcloned by limiting dilution to obtain stable monoclonal cell lines. Rat antibody clones RTNA 12E6 (IgG2c) (IP), RTNA 5F6, and RTNA 12E6 (IgG2c) (WB) and RTNC 17F1 (IgG2a) (IF) were used in this work.

**Protein immunoprecipitation**

IPSC-derived neural stem cells at d15 were lysed in ice-cold lysis buffer (1% NP-40 in TBS, pH 7.6, 2mM EDTA, 2 mM EGTA supplemented with 1x cOmplete Protease inhibitors (Roche)) for 30-45min on ice with trituration and vortexing. The lysates from at least 4 biological replicates of control and 4 matched mutant differentiation cultures were cleared at 15000 × g at 4°C for 20min before protein concentration was determined using the DCTM Protein Assay (Bio-Rad 5000111). In the meantime, 10µl Dynabeads® Protein A (Invitrogen 10002D) and 10µl Dynabeads® Protein G (Invitrogen 10004D) were incubated with 10µg RTTN A12E6 antibody supernatant or isotype control rat IgG2c (Biozol, IMS-RIGG2CPU-500) antibody at 4°C for 1h. After 3 washes with lysis buffer, 5mg of protein lysate was incubated end-over-end with the antibody-bound beads at 4°C for 2 h. Unspecific complexes were removed by washing 4 times with lysis buffer end-over-end before proteins were eluted by boiling protein-bead complexes at 95°C with a mixer at 800rpm for 10min in 2x Laemmli buffer. Samples were digested with LysC and trypsin for subsequent mass spectrometric measurements, applying a modified filter-aided sample preparation procedure as described[5, 6]. Eluted peptides were acidified with TFA and stored at -20°C.

**Label-free liquid chromatography tandem mass spectrometry**

Bottom-up LC–MS/MS measurement was performed on a QExactive HF-X mass spectrometer (Thermo Fisher Scientific) coupled online to a Ultimate 3000 RSLC nano-HPLC (Dionex). Peptides were trapped on the C18 pre-column (PepMap™ Neo 5 µm C18 300 µm x 5 mm; Thermo), eluted and separated on the C18 reversed-phase analytical column (nanoEase MZ HSS T3, 100 Å, 1.8 µm, 75 µm × 250 mm; Waters) in a 95-min nonlinear acetonitrile gradient from 5% to 40% at a flow rate of 250 nL/min. MS spectra were recorded at a resolution of 60,000 with an automatic gain control (AGC) target of 3 × 10E6 and a maximum injection time of 30 msec from 300 to 1500 m/z. From the MS scan, the 15 most abundant peptide ions were selected for fragmentation via HCD with a normalized collision energy of 28, an isolation window of 1.6 m/z, and a dynamic exclusion of 30 sec. MS/MS spectra were recorded at a resolution of 15,000 with an AGC target of 1 × 10E5 and a maximum injection time of 50 msec. Unassigned charges and charges of +1 and >+8 were excluded from the precursor selection.

LC-MS/MS raw files were analyzed on MaxQuant software (version 1.6.7.0 (Tyanova et al., 2016a)). The Uniprot database for Homo sapiens, including isoforms, was used (taxon identifier 9606; 75095 sequences) to identify proteins and quantify them by label-free quantification (LFQ) with a minimum ratio count of 2 and matching across replicates. Default settings were applied additionally allowing for the following variable modifications: oxidation (M) and deamidation (NQ). The reverse decoy database and thresholds set to FDR 1% (at peptide-spectrum match and at protein levels) were employed to exclude false positive hits. The files from *RTTN*-mutated cells included in the MaxQuant search and analysis were not considered further in the Perseus quantitative and statistical analysis.

Identification of bait interactions was carried out in Perseus 1.6.14.0 27 by analyzing the ProteinGroups.txt output table generated by MaxQuant. First, proteins identified only by site modification, decoy database (reverse) hits, potential contaminants, and proteins lacking unique peptides were excluded from the dataset using the software settings, followed by manual removal of keratins and immunoglobulins. LFQ intensities were $\log_2$-transformed before proteins with fewer than 3 valid values in at least one group (control or RTTN IP) were filtered out, and missing values were imputed based on a normal distribution. Enrichment of proteins in the IP group against the negative control group was tested by an unpaired one-tailed Student's t-test. The resulting data were visualized on volcano plots with a fold change of 1.5 and $p<0.05$, and candidates were selected if overlapping with those significant by Welch's t-test (s0.6, $p<0.05$).

**Single-cell transcriptomic analysis**

Publicly available single-cell RNA-sequencing datasets from first-trimester human brain development were analyzed to examine RTTN expression across developmental lineages and cell-cycle states. Processed expression matrices and metadata from Braun et al. (2023)[1].

Single-cell analyses and UMAP visualizations were based on data and metadata from Braun et al. (2023)[1]. Gene expression values were clipped at the 99th percentile for visualization purposes. UMAP coordinates were taken directly from the published dataset. All analyses and visualizations were performed in Python (v3.12) using Scanpy.

RTTN expression levels were quantified and analyzed using R. across annotated neural and glial populations, developmental stages, and cell-cycle phases. Statistical comparisons between cell populations were performed using Wilcoxon rank-sum tests with Benjamini-Hochberg correction for multiple testing. Heatmaps and dot plots were generated using normalized log-transformed expression values.

**RTTN expression RNA-Seq**

Bioinformatic analysis of *RTTN* mRNA expression from postmortem brain samples showed that this gene is expressed at higher levels during embryonic development (post-conception weeks, PCW, 7- 37) than its levels in brain samples of individuals with ages ranging from three months to 40 years (n=237, 287, prenatal and postnatal stages, respectively), BrainSpan Atlas[7].

**Brain-on-chip organoids**

Brain organoids were generated and cultured using a microfabricated on-chip platform adapted from Karzbrun et al.[8] Human pluripotent stem cells were dissociated and seeded into ultra-low-attachment V-bottom 96-well plates at a density of 900 cells per well in naïve medium supplemented with a ROCK inhibitor (Medchem Express HY-10583) to promote aggregation. Embryoid bodies formed within 24–36 h, after which neuronal induction medium was added, and cultures were maintained for an additional 48 h.

Microfabricated culture devices were generated by drilling 1.5-mm holes into 6-cm tissue culture dishes, followed by attaching a semi-permeable polycarbonate membrane with a UV-curable adhesive. Coverslip spacers (150 µm thickness) were fabricated using PDMS molds and NOA adhesive to define the chamber height.

On day 4, individual embryoid bodies were transferred onto membrane-covered holes and enclosed by placement of the coverslip spacer, thereby forming a sealed chamber between the membrane and coverslip. Devices were UV-cured to ensure sealing while minimizing organoid exposure. Neuronal induction medium was added above the membrane, allowing passive diffusion into the chamber. Devices were maintained at 37 °C and 5% CO2 with medium exchanges every other day.

At day 8, organoids were embedded in Matrigel introduced through inlet ports while devices were maintained on ice, followed by gelation at 37 °C. Thereafter, neuronal differentiation medium supplemented with EGF (20 ng/ml; Peprotech AF-100-15) and FGF2 (20 ng/ml; Protein Production Unit, Weizmann Institute of Science) was used to support tissue growth and maturation.

This confined culture system restricts growth in the z-dimension while allowing lateral expansion, enabling efficient nutrient diffusion and long-term imaging-compatible culture. The thin chamber geometry facilitates high-resolution real-time imaging using widefield or confocal microscopy without sectioning.

**Hippocampal organoids**

Hippocampal organoids were generated according to the Sasi protocol[9]. Human embryonic stem cells were maintained on Matrigel-coated plates or mouse embryonic fibroblasts (MEFs) in naïve human stem cell medium supplemented with 10 µM ROCK inhibitor (Medchem express HY-10583) for 24 h before aggregation. Cells were cultured to near confluency prior to aggregation, and the day of aggregation was defined as day 0.

For aggregation, cells were dissociated using trypsin for 4 min at 37°C, triturated to generate a single-cell suspension, and transferred into serum-containing medium to neutralize trypsin. Cells were seeded into low-adhesion V-bottom 96-well plates at a density of 9,000 cells per well in 100 µl differentiation medium supplemented with 20 µM ROCK inhibitor (Medchem express HY-10583) under 5% CO2.

Differentiation medium consisted of Dulbecco's Modified Eagle Medium (DMEM) (Gibco 11965092) supplemented with 20% KnockOut™ Serum Replacement (KSR) (Gibco 10828028) 0.1 mM MEM Non-Essential Amino Acids (NEAA), 100X (Gibco 11140050), 1 mM sodium pyruvate, 0.1 mM 2-mercaptoethanol, and penicillin/streptomycin. From day 0 to day 18, cultures were supplemented with 3 μM IWR1C and 5 μM SB431542. Between days 18 and 21, aggregates were transferred to 10-cm plates and cultured in DMEM/F12 supplemented with 1% GlutaMAX, 1% N2 supplement, 1% chemically defined lipid concentrate, 10% fetal bovine serum, and penicillin/streptomycin, and supplemented with 3 μM CHIR99021 and 0.5 nM BMP4.

From day 21 to day 28, organoids were maintained in the same medium without CHIR99021 or BMP4. From day 28 onward, cultures were transitioned to Neurobasal medium supplemented with B27 without vitamin A, 1% L-glutamine, penicillin/streptomycin, and 10% fetal bovine serum for continued maturation.

**Telencephalon organoids**

Human embryonic stem cells were maintained in a naïve state in HENSM medium[10] on MEFs and adapted to feeder-free conditions on Matrigel in mTeSR or StemFlex medium for one passage prior to differentiation.

For organoid formation, hESCs were dissociated into single cells and seeded at 1.5 × $10^{\wedge 5}$ cells per well in AggreWell 800 plates in mTeSR or StemFlex medium supplemented with 10 μM Y-27632. Plates were centrifuged at 500 × g for 3 min and incubated at 37 °C and 5% CO2.

After 48 h, neural induction was initiated by gradual replacement with neural induction medium (NIM; DMEM/F12 supplemented with N2, GlutaMAX, and NEAA) containing 100 nM LDN-193189, 10 μM SB431542, and 2 μM XAV-939. By day 7, aggregates were transferred to untreated plates and cultured on orbital shakers at ≤40 rpm.

Telencephalic identity was induced by supplementation with 150 ng/ml FGF8 from day 10 to day 15. From day 16 to day 30, organoids were maintained in NIM, with 10 ng/ml hLIF added starting on day 20. At day 30, organoids were transferred to long-term survival medium consisting of Neurobasal supplemented with N2, B27, GlutaMAX, NEAA, sodium pyruvate, 1% KSR, and 0.1% Matrigel and cultured under 40% oxygen conditions on orbital shakers.

From day 50 onward, neuronal maturation was enhanced using GENtonik supplementation containing 1 μM GSK2879552, 1 μM EPZ-5676, 1 μM NMDA, and 1 μM Bay K8644. After day 60, B27 supplemented with vitamin A was used to support further maturation.

**O-propargyl-puromycin (OPP) protein synthesis assay**

Protein synthesis was measured using an O-propargyl-puromycin (OPP; Vector Laboratories, Cat# CCT-1407-5) incorporation assay [11]. hESCs were plated on Matrigel-coated Ibidi plates (μ-Slide 18 Well 81816) and cultured until well-defined colonies formed. For hippocampal organoid experiments, experiments were repeated on days 53, 80, 84, 90, 100, 120, and 125. At least three organoids per condition were dissociated using TrypLE Express for 10 min, plated onto Ibidi plates, and allowed to recover prior to the assay.

OPP was diluted 1:1000 in culture medium to a final concentration of 20 µM, and cells were incubated with 100 µl OPP-containing medium per well for 60 min under standard culture conditions. Following incubation, cells were washed once with PBS and fixed in 3.7% formaldehyde in PBS for 15 min at room temperature. Cells were then permeabilized with 0.5% Triton X-100 in PBS for 15 min, followed by two washes with PBS.

A click-reaction mixture was freshly prepared in PBS containing 100 mM Tris (pH 8.5), 1 mM $CuSO_4$, 2.5 µM azide-conjugated fluorophore, and 100 mM ascorbic acid (added immediately before use). Cells were incubated with 100 µl reaction mixture per well for 30 min at room temperature, protected from light. Following the reaction, cells were washed three times with PBS. Nuclei were stained with Hoechst 33342 (1:2000 in PBS), and samples were subsequently imaged and analyzed.

Cells were imaged using a Dragonfly spinning-disk confocal microscope and analyzed with Imaris software to detect and quantify OPP-positive spots. At least three independent biological replicates were analyzed for each genotype.

**Brain Organoid Polysome profiling**

**Polysome profiling assay**

Polysome profiling was performed to assess global translation, ribosome biogenesis, and translation elongation in telencephalon and hippocampal organoids. All procedures were carried out under RNase-free conditions using RNase Zap-treated surfaces and equipment. Standard polysome profiling buffer consisted of 20 mM Tris-HCl (pH 7.5), 150 mM NaCl, 5 mM $MgCl^2$, 100 µg/mL cycloheximide (CHX), and 1 mM DTT. For sucrose gradient preparation, 10–50% sucrose solutions were prepared in polysome buffer, and gradients were generated in open-top polyclear centrifuge tubes (14 × 89 mm, Seton) using a Gradient Master (Biocomp). Gradients were stabilized overnight at 4°C prior to use.

Telenchephelon organoids at day 30 and day 60 were treated with 100 µg/mL CHX and washed with ice-cold PBS containing CHX. Organoids were lysed in polysome lysis buffer containing 1% Triton X-100, Turbo DNase, and RiboLock RNase inhibitor in polysome gradient buffer. Lysates were sonicated on ice using a Branson Sonifier 250 and centrifuged at 12,000 × g for 10 min at 4°C to remove debris. Equal amounts of lysate, normalized by OD 260, were carefully layered onto sucrose gradients. Gradients were centrifuged at 35,000 rpm for 3 hours at 4°C in an ultracentrifuge using an SW41 rotor.

Following centrifugation, gradients were fractionated using a Biocomp Gradient Fractionator coupled to a Triax flow cell, and 21 500 µl fractions were collected for downstream analyses. Fractions were either processed immediately or stored at −80°C. For half-mer analysis, hippocampal organoids at day 50 were treated with 100 µg/ml CHX for 10 min prior to lysis.

For ribosome biogenesis assays, sucrose gradients and lysis buffers were prepared without $MgCl^2$ and CHX and supplemented with 30 mM EDTA. EDTA chelates $Mg^{2+}$ ions required for ribosome stability, resulting in dissociation of polysomes and 80S ribosomes into individual 40S and 60S ribosomal subunits. This approach enabled assessment of ribosome assembly and subunit distribution as a measure of ribosome biogenesis defects.

**Polysome-Seq**

RNA was purified from pooled fractions corresponding to the 80S monosome peak, light polysomes (LP, mRNAs associated with 2-4 ribosomes), and heavy polysomes (HP, ≥5 ribosomes) using the RNA Clean & Concentrator kit (Zymo Research), including on-column DNase digestion. RNA was eluted in RNase-free water and quantified by spectrophotometry.

For RNA sequencing, 150 ng of total RNA or polysome fraction-derived RNA per sample was used for library preparation with the TruSeq Stranded Total RNA Library Prep kit (Illumina; Human/Mouse/Rat). Libraries were sequenced on a NovaSeq X platform to a depth of ~50 million reads per sample.

Adapter trimming was performed using Cutadapt, followed by alignment to the hg38 human reference genome and generation of a gene-level count matrix using STAR v2.7. These steps were carried out using the UTAP2 pipeline[12]. Normalization of the count matrix and differential expression analysis were performed using DESeq2 (v1.36.0)[13]. The statistical model included genotype, polysome fraction, and their interaction term. Genes with a baseMean > 10, |log2 fold change| > 0.6, and adjusted p-value (padj) < 0.05 were considered to exhibit significant changes in polysome-associated RNA levels in the mutant compared to the control.

Translational efficiency analyses were performed by comparing fraction-enriched transcript abundance relative to total RNA abundance:

$$TE = log_2\left(\frac{HP_{mut}}{HP_{ctrl}}\right) - log_2\left(\frac{Total_{mut}}{Total_{ctrl}}\right)$$

Gene ontology enrichment analyses were performed using ShinyGO 0.85.1 analysis[14].

Codon usage frequencies and transcript length distributions were calculated using custom R scripts implemented with the Biostrings package (v2.72.1). Calculations were performed using Ensembl canonical transcripts, with a single transcript retained per gene.

For the codon abundance analysis, the terminal stop codons (TAA, TAG, and TGA) were removed prior to analysis, and codons containing ambiguous nucleotides were excluded. Codon counts were tallied across all 64 possible codons, including those absent from the dataset (assigned a count of zero). Codon usage frequencies from enriched and depleted gene sets were compared against global background codon frequencies, and enrichment significance was assessed using Fisher's exact test followed by the Benjamini–Hochberg false discovery rate (FDR) correction. Codons with p.adj < 0.05 and |log2FC| > 1 were considered significantly enriched or depleted.

**Electron microscopy**

Hippocampal organoids at day 90 were fixed overnight with a solution of 2.5% glutaraldehyde, 4% paraformaldehyde in 0.1 M cacodylate buffer. They were then rinsed with 0.1 M cacodylate buffer containing 5 mM CaCl2, and then fixed and stained for one hour with an osmium tetroxide solution (1% OsO4, 0.5% potassium dichromate, 0.5% potassium hexacyanoferrate in 0.1 M cacodylate buffer), rinsed with 0.1 M cacodylate buffer followed by milli-Q H2O, and then stained for another hour in 2% uranyl acetate. They were rinsed again with Milli-Q $H_2O$, dehydrated with a series of ethanol dilutions (50%, 70%, 96%, 100%), and then transferred to 100% acetone. Organoids were then embedded in epon (EMS, PA, USA) diluted with anhydrous acetone at 25%, 50%, 75%, and 100%, and finally embedded in 100% epon within rectangular molds, dried at 60°C for 24 hours, and stored until sectioning. Sample blocks were sectioned to a thickness of 70 nm with an EM UC7 ultramicrotome (Leica microsystems, Vienna, Austria) equipped with an Ultra 45° diamond knife (Diatome Ltd, Nidau, Switzerland) and were mounted on either formvar and copper-coated 200 mesh copper grids (EMS) or formvar - coated 2x1 mm copper slot grids (EMS). Sections were stained with Reynolds lead citrate and imaged at room temperature using three different microscopes: 1. a Tecnai T12 transmission electron microscope (Thermo Fisher Scientific, The Netherlands) at 120 kV equipped with a bottom-mounted TVIPS TemCam-XF416 4k × 4k CMOS camera using TVIPS- EMplified software, 2. a FEI Tecnai G2 F20 transmission electron microscope (Thermo Fisher Scientific, The Netherlands) at 200 kV equipped with a bottom-mounted TVIPS TemCam-XF416 retractable 16-megapixel CMOS camera using TVIPS- EMplified software, and 3. a Talos Arctica TEM microscope (Thermo Fisher Scientific, The Netherlands) at 200 kV equipped with a Gatan OneView camera (Gatan Inc., California, USA) .

**TEM image analysis**

For each mutant and control, three independent organoids were analyzed, and 5-14 micrographs per organoid (from one or two sections). Images were analyzed using the ParticleSizer plugin in Fiji (NanoDefine). Ribosome density was quantified by calculating the number of ribosomes per µm² within defined regions of interest (ROIs). Statistical analysis was performed using a nested one-way ANOVA, followed by Holm-Šídák's multiple comparisons test.

**Protocol for BACS treatment and tRNA library preparation**

Total RNA was first deacylated by brief alkaline treatment (Tris-HCl, pH 9.0, 37°C) to remove aminoacyl groups from the tRNA 3′ end, followed by magnetic bead-based (silane) purification. The 3′ ends of the RNA were then enzymatically dephosphorylated (FastAP) and treated with DNase (Turbo DNase) together with T4 polynucleotide kinase in a combined reaction to generate ligatable 3′-hydroxyl ends, again followed by bead-based cleanup. A barcoded RNA adapter was ligated to the 3′ end of each sample using T4 RNA ligase in a high-percentage PEG/DMSO buffer system to improve ligation efficiency, allowing individual samples to be uniquely indexed and subsequently pooled. Pooled, barcoded samples were size-selected for small RNAs in the tRNA size range by column-based purification, discarding the long-RNA fraction. BACS treatment was then performed as describe[15] to enable base-resolution detection of pseudouridine prior to reverse transcription. Reverse transcription of the adapter-ligated, pooled, size-selected, and BACs-treated tRNA pool was carried out using a primer complementary to the ligated 3′ adapter and Maxima reverse transcriptase. Residual RT primer and unincorporated oligonucleotides were removed enzymatically (ExoSAP-IT), and the RNA template strand was then hydrolyzed by brief alkaline treatment (NaOH, 70°C) and neutralized (HCl), leaving single-stranded cDNA. A second 5′-end adapter was ligated to the cDNA (again via T4 RNA ligase in PEG/DMSO buffer), completing a sequencing-ready construct flanked by defined adapter sequences at both ends. The resulting cDNA was PCR-amplified using indexed primers and a high-fidelity polymerase (KAPA HiFi) under a touchdown-style cycling program, an initial low-stringency annealing phase followed by additional higher-temperature amplification cycles, to generate the final sequencing library. Libraries were assessed by gel electrophoresis to confirm the expected size distribution (predominantly 200-300 bp, distinct from primer-dimer products near 140 bp), cleaned up by SPRI bead-based size selection to remove residual adapter dimers, and quantified by qubit assay prior to pooling for sequencing.

**Sequencing data metrics**

Libraries were sequenced on an Illumina NovaSeq X instrument using a paired-end, dual-indexed 61-8-8-61 cycle configuration (61 bp Read 1, 8 bp Index 1, 8 bp Index 2, 61 bp Read 2), generating approximately 10 million paired-end reads per sample. Reads were adapter-trimmed using cutadapt retaining >= 30% of reads. Sequencing quality was assessed using FastQC(FastQC/0.12.1-Java-11) and summarized with MultiQC(MultiQC/1.12-foss-2021b), demonstrating consistently high-quality reads, with mean per-read Phred quality scores of approximately Q39-Q40 across all libraries.

**Detailed description of the tRNA sequencing and data-analysis pipeline**

Raw sequencing reads were first quality- and adapter-trimmed from FASTQ files to remove residual adapter sequence and low-quality bases. Read 2 reads were clustered and aligned to the human hg38 tRNA reference the mim-tRNAseq pipeline[16], which performs cluster-aware alignment across the highly redundant tRNA gene set. Read 1 alignments were of poor quality, whereas Read 2 yielded reasonable, high-confidence unique alignments. Since BACS relies on detecting misincorporation events at individual positions rather than full-length transcript coverage, only Read 2 reads were carried forward for alignment and all downstream analysis. The resulting single-end (Read 2) alignments (BAM files) from mimseq pipeline were processed with txtools[17], an R package that integrates genome alignments with transcriptomic coordinates to generate single-nucleotide-resolution pileup metrics, including per-base coverage, nucleotide identity, and mismatch counts; txtools natively supports single-end, Read 2-only input via dedicated processing arguments. This single-nucleotide-resolution output was used to quantify, at each candidate T-reference position, the BACS-induced T-to-C conversion signature diagnostic of pseudouridine, yielding the raw conversion per-site, per-sample counts and conversion fractions underlying all downstream statistical analyses (see Statistical methods and significance criteria, above) and figures.

**Software, packages, and versions used for the analyses**

| **Category** | **Tool / Package** | **Version** | **Used for** |
|---|---|---|---|
| Sequencing / alignment | cutadapt | 4.2 | Adapter trimming |
| Sequencing / alignment | STAR | 2.7.9a | rRNA-filtering alignment (QC) |
| Sequencing / alignment | SAMtools | 1.21 | BAM file processing |
| Sequencing / alignment | Singularity | 3.8.0 | Container runtime for the mim-tRNAseq container |
| Sequencing / alignment | mim-tRNAseq | 1.3.11 | tRNA clustering and alignment |

| | | | |
|---|---|---|---|
| Sequencing / alignment | txtools | 1.0.6 | Single-nucleotide-resolution pileup extraction (BACS T-to-C conversion) |
| R environment | R | 4.2.3 | Statistical analysis and figure generation |
| R packages | dplyr | 1.1.4 | Data wrangling |
| R packages | tidyr | 1.3.1 | Data wrangling |
| R packages | purrr | 1.0.2 | Functional iteration |
| R packages | stringr | 1.5.1 | String handling |
| R packages | forcats | 1.0.0 | Factor handling |
| R packages | ggplot2 | 3.5.1 | Plotting |
| R packages | scales | 1.3.0 | Plot scaling / axis formatting |
| R packages | knitr | 1.47 | Report rendering |
| R packages | rmarkdown | 2.27 | Report rendering |
| R packages | systemfonts | 1.3.2 | Font handling |
| R packages | ragg | 1.2.5 | Raster graphics device |
| R packages | svglite | 2.2.2 | SVG export |
| R packages | gridExtra | 2.3 | Multi-panel plot layout/composition |
| R packages | gtable | 0.3.5 | Grid-based table/plot layout |
| R packages | data.table | 1.15.2 | Fast data handling |

**Statistical methods and significance criteria**
Two statistical frameworks were used, each matched to its own data source and purpose:

***Framework 1: group-mean comparison (heatmap figure, Panel d).*** Per-sample Ψ (BACS T→C conversion fraction) at each candidate T-reference position was parsed from the aggregate conversion table (merged_conversion.csv; n = 4 biological replicates per condition: Control, $PUS7L^{KO7}$, $PUS7L^{KO66}$, $RTTN^{25}$, $RTTN^{C27Y}$). Candidate positions were restricted to those with a maximum group mean Ψ > 5% (480 sites tested). Each KO condition was compared to Control using a two-sided Welch's t-test (unequal variances), with Benjamini-Hochberg FDR correction applied within each KO-vs-Control comparison. Significance is reported as *q < 0.05, **q < 0.01, ***q < 0.001.

***Framework 2: per-replicate GLM (volcano figure, Panel c).*** Raw per-position (C, T) read counts were recovered from the per-replicate alignment count files (20 libraries: 5 conditions × 4 replicates), restricted to the same tRNA gene panel and candidate T-positions, with an additional depth-based QC filter (minimum group mean read depth ≥ 10) not applied in Framework 1. This yielded 332 candidate sites. For each site and each KO/Rotatin mutant-vs-Control comparison, a quasi-binomial GLM (cbind(C, T) ~ group) was fit and compared against a null (intercept-only) model by an F-test; resulting p-values were BH-corrected within each comparison. A site was called significant if $q < 0.05$ and $|\Delta \Psi$ (KO − Control)$| > 0.10$, a combined statistical- and effect-size threshold, applied to avoid depth-driven trivial significance at very small effect sizes.

The 11-site core PUS7L-dependent signature was defined, independently of either statistical test above, as the fixed set of sites lost in both PUS7L-KO lines and RTTN mutants ($RTTN^{25}$, $RTTN^{C27Y}$); it is used identically across all figures so that the same 11 sites can be tracked across both statistical frameworks and all five KO-mut/control comparisons.

**tRNA secondary structure visualization**

The mature human **tRNA-Leu-TAG-3-1** sequence was obtained from the Genomic tRNA Database (GtRNAdb;

https://gtrnadb.ucsc.edu/genomes/eukaryota/Hsapi38/genes/tRNA-Leu-TAG-3-1.html).

tRNA-Ala-TGC-5-1

https://gtrnadb.ucsc.edu/genomes/eukaryota/Hsapi38/genes/tRNA-Ala-TGC-5-1.html

The secondary structure was generated using the RNAcentral RNA 2D Templates (R2DT) server (https://rnacentral.org/r2dt), which predicts RNA secondary structures based on standardized template layouts. The resulting R2DT structure was subsequently imported into RNAcanvas (https://rnacanvas.app/) for manual editing, annotation, and visualization of pseudouridine sites and structural features.

**Immunostaining and confocal imaging**

Immunohistochemistry (IHC) was performed on day 65 hippocampal organoids derived from two independent batches comprising 3 and 4 organoids, respectively. Organoids were fixed in 4% paraformaldehyde for 3-4 hours at 4°C with gentle shaking, followed by overnight incubation in 20% sucrose at 4°C. Organoids were embedded in OCT, cryo-sectioned at 14 μm, and mounted onto positively charged microscope slides. Each organoid was represented four times per slide at sequential depths, and control and *RTTN* mutant organoids were mounted on the same slides. Slides were air-dried overnight prior to staining. Antigen retrieval was performed in 10 mM sodium citrate buffer (pH 6.0) at 90-95°C for 5-6 min.

Immunohistochemistry (IHC) for human telencephalic organoids were performed on day 30 and day 60. Organoids were washed with PBS once and fixed with 4% formaldehyde in PBS for 24 h at 4° C. Fixed organoids were twice transferred to 30% sucrose in PBS for 24 h each time and then embedded in Tissue-Tek OCT compound and stored at -20° until use. Embedded organoids were sliced into 20 μm sections in a Leica CM3050S cryostat and immediately transferred to SuperforstPlus adhesion microscope slides. Excess OCT was removed from the slides with PBS and antigen retrieval was performed with 10 mM citrate buffer boiling for 8 min. Slides were blocked with 10 % Horse serum, 10 % FBS, and 0.1 % TritonX-100 for 1 h.

For monolayer cell immunostaining, cells were plated on polyornithine/laminin-coated dishes and fixed with either 4% paraformaldehyde for 10 min at room temperature or 100% methanol for 10 min at −20°C for centrosomal staining experiments. Samples were blocked in 10% horse or goat serum supplemented with 0.1–0.5% Triton X-100 in PBS for 1 h at room temperature and incubated with primary antibodies overnight at 4°C. Alexa Fluorophore-conjugated secondary antibodies (Life Technologies or Jackson ImmunoResearch) were applied for 1–2 h at room temperature together with Hoechst 33342 or DAPI nuclear staining. Coverslips were mounted using Aqua Polymount (Polysciences).

Primary antibodies used included ACINUS (Acin1) (rabbit, 1:1000, Abcam), Arl13b (rabbit, 1:500, Proteintech), RPL11 (rabbit, ab79352 abcam), KI67 (rabbit, 1:100, Millipore), pHH3 (rabbit, 1:100, Millipore), cleaved Caspase-3 (rabbit, 1:200, Cell Signaling), CDK5RAP2 (rabbit, 1:1000, Sigma Aldrich), FOXG1 (rabbit, 1:100, Abcam), GFAP (rabbit, 1:2000, DAKO), NESTIN (mouse 10c2, 1:400, Millipore), MAP2 (mouse, 1:500, Sigma Aldrich), MATR3 (rabbit, 1:500, Abcam), NeuN (mouse, 1:200, Millipore), Ninein (rabbit, 1:200, Bethyl), PAX6 (rabbit, 1:100, Covance), PAX6 (rabbit, 1:300, Biolegend), PRPF6 (mouse, 1:500, Santa Cruz), SMI312 (mouse, 1:200, Covance), SOX2 (mouse, 1:200, Santa Cruz), TUJ1 (guinea pig, 1:250, SYSY), HOPX (rabbit, 1:200, Sigma-Aldrich), LEF1 (rabbit, 1:200, Cell Signaling), VIRMA (Kiaa1429) (rabbit, 1:1000, Proteintech), ZBTB20 (rabbit, 1:200, Sigma-Aldrich), pVIM (mouse, D076-3), OTX2 (rabbit, 13497-1-AP), FOXG1 (rabbit ,AB196868), CTIP2 (rat, AB18465),

Confocal images were acquired using a Dragonfly spinning-disk confocal microscope with 20×, 40×, or 63× objectives or an Olympus FV1000 confocal laser-scanning microscope using 20X/0.85 N.A. or 40X/1.35 N. An objective or on a Zeiss LSM710 with a 40X/1.1 N.A. objective in 0.5-1μm z-stack steps. Image processing and analysis were performed using Imaris, ImageJ, and Adobe Illustrator/Photoshop by applying equal brightness/contrast adjustments to control and mutant cell images.

**Western blot**

Dorsal forebrain cells at d15 of differentiation were collected, lysed in ice-cold lysis buffer and the lysates cleared as done for immunoprecipitation. After determination of protein concentration, 100ug were loaded on 6% poly-acrylamide SDS gels and transferred to 0.22 μm nitrocellulose membrane. After transfer, membranes were washed in TBST (TBS with 0.1% Tween-20, pH7.4), blocked with 5% non-fat dry milk in TBST for at least 1h at room temperature (RT), incubated overnight with primary antibodies in 1% milk in TBST and on the next day, with secondary HRP-conjugated secondary antibodies in 1% milk in TBST for 1h at RT. The signal was detected with ECL reagents.

.

**RT-qPCR analysis of rRNA biogenesis processing**

To assess rRNA processing intermediates, total RNA was isolated from hippocampal organoids at day 50 using the Quick-RNA™ Miniprep Plus Kit (Zymo Research, R1058). For each biological replicate, lysates were generated by pooling 8 organoids, and four independent biological replicates were analyzed per genotype. cDNA synthesis was performed using the High-Capacity cDNA Reverse Transcription Kit (Applied Biosystems, 4368814). Quantitative PCR assays targeting the 5′ ETS, ITS1, ITS2, 18S, 5.8S, 28S, and 3′ ETS regions of the human rDNA transcription unit were performed using Fast SYBR™ Green Master Mix (Applied Biosystems, 4385612). Quantitative PCR was carried out using a StepOnePlus™ Real-Time PCR System (Applied Biosystems). Expression values were normalized to UBC using the $2^{-\Delta Ct}$ method, and the relative abundance of rRNA processing intermediates was compared across genotypes.

| | |
|---|---|
| ITS1-F | GAAACCTTCCGACCCCTCTC |
| ITS1-R | GAGGCCCTTCCTGGCG |
| ITS2-F | TCCGGGTTCCTCCCTCG |
| ITS2-R | GCACGGGACCTTCCACC |
| 3' ETS-F | CTTCTTCGGTTCCCGCCTC |
| 3' ETS-R | AACCACGCTCCCCGGAC |
| 5' ETS-F | TCTGTGCCCTCTTCCCCG |
| 5' ETS-R | ACCAACGGACGTGAAGCC |
| 5.8S rRNA-F | GTGCGTCGATGAAGAACGC |
| 5.8S rRNA-R | AGTGCGTTCGAAGTGTCGAT |
| 28S rRNA-F | ATCAGACGTGGCGACCCG |
| 28S rRNA-R | CTGTTCACTCGCCGTTACTG |
| 18S rRNA-F | CGGCTACCACATCCAAGGAA |
| 18S rRNA-R | GCTGGAATTACCGCGGCT |

| | |
|---|---|
| 5S rRNA-F | GCCCGATCTCGTCTGATCTC |
| 5S rRNA-R | AGCCTACAGCACCCGGTATT |
| UBC F | CGTCGCAGCCGGGATT |
| UBC R | ATCTGCATTGTCAAGTGACGA |

**5-Ethynyl Uridine (5-EU) incorporation assay**

Nascent RNA synthesis was measured using 5-ethynyl uridine (5-EU, click chemistry tools #1261-10) incorporation[18]. Dissociated organoids at day 76 and day 97 were incubated with 200 μM 5-EU for 1 h at 37 °C, followed by fixation and click-chemistry detection using fluorescent azide reagents. Images were acquired by confocal microscopy, and fluorescence intensity measurements were quantified using Imaris.

**Live imaging and mitotic progression analysis**

Brain-on-chip organoids expressing membrane-targeted LYN-GFP and H2B-mCherry were imaged by time-lapse confocal microscopy under standard cell culture conditions to monitor mitotic progression and interkinetic nuclear migration. Live imaging was performed on organoids at two developmental stages (days 13-14 and 20-22), and a total of 4-6 independent live-imaging movies were analyzed, with each movie considered an independent experimental batch. Individual mitotic cells were manually tracked through prometaphase, metaphase, anaphase, and telophase to determine total mitotic duration and the length of each mitotic phase. Prometaphase, metaphase, and anaphase durations were quantified separately. Statistical analysis was done using Student's t-test (2-tailed).

**Interkinetic nuclear motility**

Interkinetic nuclear motility (IKNM) was analyzed using time-lapse imaging of brain-on-chip organoids using a Dragonfly spinning-disk confocal microscope under standard culture conditions. Live imaging was performed on day 14 and day 20 organoids, with data collected from 3-4 independent live-imaging movies, each representing an independent experimental batch. Time-lapse z-stack images were acquired at defined intervals and processed using Imaris software. Individual nuclei within the ventricular zone-like regions were manually tracked over time using the Imaris tracking module. Nuclear trajectories were used to quantify migration velocities during both apical and basal movements along the apical-basal axis.

**EdU incorporation assay**

DNA synthesis was assessed using 5-ethynyl-2′-deoxyuridine (EdU) incorporation. Dissociated organoids were incubated for 30 minutes with 10 μM EdU prior to fixation and processed using click-chemistry detection[18]. The percentage of EdU-positive nuclei was quantified across multiple dissociated organoids and imaging fields.

**RNA-sequencing and alternative splicing analysis**

Bulk RNA-sequencing libraries were generated from hippocampal organoids at day 49 using total RNA. Libraries were sequenced on Illumina platforms to generate paired-end reads.

**RNAseq analysis**

Raw RNA-seq data were processed using the UTAP pipeline[19], which includes adapter trimming with Cutadapt (parameters: --a ADAPTER1 -a "A{10}" -a "T{10}" -A "A{10}" -A "T{10}" --times 2 -q 20 -m 25), alignment to the human reference genome (hg38) using STAR v2.4.2a (parameters: --alignEndsType EndToEnd, --outFilterMismatchNoverLmax 0.05), read quantification against GENCODE v34 gene annotations, and differential expression analysis using DESeq2[13].

Normalization and differential expression analyses were performed using DESeq2 with betaPrior = TRUE, cooksCutoff = FALSE, and independentFiltering = FALSE. Raw p-values were adjusted for multiple testing using the Benjamini-Hochberg procedure. Differentially expressed genes were defined by an absolute $\log_2$ fold change > 1, an adjusted p-value < 0.05, and a mean base count > 10.

Alternative splicing analysis was performed using MAJIQ v2.2[20]. Splice graphs were constructed using MAJIQ against GENCODE v36 primary assembly annotations (GRCh38), applied to pairwise comparisons of every mutant versus wild-type, in each organoid day. Local splicing variations (LSVs) were considered differentially spliced if the expected |ΔPSI| exceeded [0.15] with a posterior probability ≥ [0.95], in accordance with MAJIQ default parameters. Gene ontology enrichment analyses of differentially spliced genes were performed using clusterProfiler.

**Nucleofection and time-lapse imaging of neural cells**

Time-lapse imaging was performed at 37°C with 5% $CO_2$ on an Olympus FV1000 inverted confocal laser-scanning microscope using a 20X/0.7 N.A. objective and a humidified stage-top incubator (Tokai Hit). Confocal image stacks with a 0.5-1µm step size were collected every 20min for 16-72h, and quantifications were done on movies generated from maximum projection in FIJI/ImageJ.

**Statistical analysis**

Statistical analyses were performed using GraphPad Prism, R, and Microsoft Excel. Normality of datasets was assessed prior to statistical testing. Two-group comparisons were performed using two-tailed Student’s t-tests or Mann-Whitney U tests, as appropriate. Multiple-group comparisons were analyzed using one-way ANOVA, Welch's ANOVA, Kruskal-Wallis tests, or nested analyses followed by appropriate post hoc multiple-comparison corrections, as indicated. For RNA-sequencing analyses, multiple-testing correction was performed using the Benjamini-Hochberg method. Statistical significance was defined as adjusted $P < 0.05$ unless otherwise indicated. Exact statistical tests, sample sizes, and P values are provided in the corresponding figure legends.

Unless otherwise indicated, at least three independent biological replicates were analyzed for each experiment or, in the case of iPSC-derived models, at least three independent differentiation experiments including distinct patient-derived clones. Data are presented as mean or median ± interquartile range (IQR).

**Data availability**

The mass spectrometry proteomics data have been deposited in the ProteomeXchange Consortium via the PRIDE[21] partner repository, with the dataset identifier PXD077974. RNA-sequencing and polysome-sequencing datasets generated in this study were deposited in a public repository, GEO, accession numbers GSE337409 and GSE337408.
